# CRISPR/Cas9 Optimization Using Electronic Genome Mapping: Potential Role for Studying Human Genetic Disease

**DOI:** 10.64898/2026.09.14.751601

**Authors:** Lindsay Opsahl, Wentao Huang, Syndi Koltz, Kaylee L. Mathews, John F. Thompson

**Affiliations:** Nabsys 2.0 LLC; Providence, RI USA

**Keywords:** CRISPR, genome mapping, electronic genome mapping, DNA mismatch, Protein-protein interactions, gRNA, off-target

## Abstract

CRISPR Cas9’s ability to bind targeted and non-targeted sites and its functional interactions with other proteins and domains have been studied using many technologies. These studies are often limited by using oligos as substrates or by a reliance on predicted binding sites. We show how electronic genome mapping (EGM) with its ability to detect single-molecule DNA nicks can determine how gRNA mismatches impact targeted versus off-target binding across the whole human genome *in vitro*. Cas9 was found to be capable of binding a single site adjacent to the *FXN* repeat expansion responsible for Friedreich Ataxia using a gRNA specific for that sequence while gRNAs matching a repetitive sequence were able to bind hundreds of sites. Binding to perfectly matched gRNA sites was compared to sites with mismatches at each gRNA position. The PAM site and nearby seed region bases were confirmed as critical for binding. In contrast, single mismatches toward the gRNA 5’ end allowed off-target binding. We also used EGM to determine how physically close the bound dCas9/gRNA complex can be relative to nearby DNA modification sites and still allow activity. Bound CRISPR/dCas9 inhibits nickases with recognition sites 4-7 bp away from the gRNA/PAM. Because EGM can simultaneously evaluate many mismatched off-target sites, these insights may be extended to other CRISPR systems and enable optimization of Cas proteins and gRNAs. This is the first example of EGM being used to characterize proteins and RNAs. This is also the first study demonstrating the combined application of CRISPR and EGM methodology and its potential role in the detection of specific repeat regions in the human genome that are associated with human repeat expansion disorders.

## Introduction

CRISPR-Cas9 has emerged as a revolutionary gene-editing technology with diverse research applications including disease modeling, cancer research, drug discovery and functional genomics [1]. Additionally, CRISPR mediated applications hold promise as a valuable tool for direct therapeutic interventions for genetic disorders such as sickle-cell anemia and cancer immunotherapy [2, 3]. The CRISPR system composed of Cas9 and its homologs complexed with RNAs have become one of the most studied binding and DNA modification systems [4, 5]. Their detailed structure and how the proteins bind DNA and RNA have been shown in detail [6]. Mutations in Cas proteins and Cas fusions with other functional domains have made CRISPR the most useful system for gene editing and introducing other DNA modifications [7–10].

The Cas9 component of CRISPR has been well characterized including mutations that eliminate double-strand cleavage. This can lead to either nicking of a single DNA strand (D10A Cas9)[11] or no cutting at all (dCas9)[12], even though the guide RNA (gRNA)/Cas9 complex still binds specifically to DNA[5, 6]. Cas9 binding is specific and long-lasting, even with mismatches in the gRNA [13–15]. The protospacer adjacent motif (PAM) sequence is not present on the 20 nt gRNA, but it has a major impact on CRISPR’s ability to bind at the appropriate locations [16]. Here, we have evaluated the ability of dCas9 to stably bind thousands of sites across the whole human genome *in vitro* in a gRNA-dependent manner and use that information to draw conclusions about activity. This information complements what has been found using other techniques like GUIDE-seq [17], DISCOVER-seq [18], CIRCLE-seq [19], and Digenome-seq [20] for identifying off-target CRISPR sites and the relative impact of gRNA mismatches on Cas9 function.

Electronic genome mapping (EGM) of ultra-long, single-molecule DNA has become an important adjunct to sequencing for characterizing long-range DNA structure, assisting genome assembly, and detecting structural variants [21, 22], but we show it can also be used to characterize protein activities. EGM is based on the same detection principle as nanopore sequencing where the physical presence of DNA or tagged DNA in a nanochannel with a voltage gradient causes a measurable current blockade [23]. This provides EGM with the advantage that DNA and site-specific tags can be detected at higher resolution and lower cost than other mapping methodologies such as optical genome mapping (OGM). While EGM, as currently configured, has an intrinsic resolution between neighboring tags on the order of hundreds of base pairs, we have designed experiments that achieve functional single-base resolution.

To generate predictable tag sites throughout the genome, the site-specific nicking enzymes Nb.BssSI and Nt.BspQI were used to introduce DNA breaks for tag attachment. The nickases have non-palindromic recognition sites of 6 and 7 base pairs, respectively. When used in combination on human genomic DNA, they make single-stranded DNA nicks approximately once every 4000 bp. This creates a broad range of intervals between sites, ranging from a few bp to more than 100,000 bp. For EGM, DNA polymerase labels the nicks with modified nucleotides. The modified nucleotides are then tagged and distance between tags is measured by recording voltage as a function of time (Supplementary Figure 1). It is generally possible to unambiguously place molecules from unique regions of the genome when a pattern of >6 interval sizes is obtained and compared to a human reference.

The programmability of the CRISPR/Cas9 system enables modification of the nickase-created tagging profiles. Treating the DNA with D10A Cas9 (nicking only) allows the introduction of new tags. Binding the DNA with the catalytically inactive dCas9 blocks nickases and prevents them from acting on the bound regions. We will show that the CRISPR/Cas9 system can block nicking at specific sequences, thereby enabling observation of interactions between Cas9 and other DNA-binding or DNA-modifying proteins at single-nucleotide resolution. Using isolated nick sites, the impact of mismatches in the gRNA can be evaluated with respect to DNA binding.

The potential of this methodology to accurately detect repeat expansions in the FXN gene, causing Friedreich ataxia (FRDA), was explored. FRDA is an autosomal recessive, progressive ataxia disorder with age of onset from early childhood (as early as two years) to early adulthood [24]. Given the severity of the presentation, it becomes important to detect the GAA repeat expansions (in ∼96% of cases with biallelic expansions) or pathogenic sequence variants in the FXN gene [25]. This is the first CRISPR/Cas9 mediated EGM study of its kind demonstrating the potential role of using this approach for the detection of a human genetic repeat expansion disorder.

## Results

While the ultimate goal is to use CRISPR to analyze human DNA, initial experiments were carried out with bacteriophage lambda DNA, providing a simple 48.5 kb model system that can be easily studied. To test the use of dCas9 for blocking Nt.BspQI and Nb.BssSI sites, sgRNAs were synthesized corresponding to distinct lambda sequences overlapping individual nick sites. Initial experiments were carried out by pre-incubating the sgRNA/dCas9 complex with DNA at 37°C for 1 hour prior to adding nickases. However, CRISPR/Cas9 has been shown to be active at temperatures as low as 4°C [13]. It was found that the pre-incubation in the absence of nickases was unnecessary when reactions were started at temperatures below the enzymatic nickase optima. sgRNA-driven dCas9 binding is rapid even on ice, and we observed no difference in behavior across varying combinations of incubation times and temperatures (0-60 minutes and 0-25°C). Other Cas proteins may have different kinetic and temperature dependencies. With dCas9, all components can be added simultaneously at 0-25°C and the mix can be raised to the temperature required for subsequent reactions. The enzymatic nickases used here, Nt.BspQI and Nb.BssSI, have optimal activity at 50°C and 37°C, respectively, with much lower activity at 20°C (https://www.neb.com/en-us/tools-and-resources/selection-charts/effect-of-various-temperatures-on-nicking-endonucleases) so do not begin nicking until after the dCas9 binds at lower temperature.

Lambda DNA was treated with the standard EGM nicking and labeling protocol which uses only Nt.BspQI as the nickase. Modified nucleotides were added to the nick sites via Klenow DNA Polymerase and unincorporated nucleotides were then removed using a Nanosep column. Modified nucleotides inserted at the nick sites were tagged and RecA-coated per the standard protocol. The resulting DNA was injected into the OhmX Analyzer and run for at least 10 minutes to collect data on more than 300,000 molecules. Based on the known nick sites for Nt.BspQI, the analysis program maps each tag to its most likely position. There is a low background level of false positive tags between true positive locations. When not blocked, site #5 has >95% true positive tags mapped while it drops to <20% tagged when the site is blocked by CRISPR. This likely underestimates the level of blocking because the nearby low-level false positives are assigned to this site as true positives when it is blocked

While the lambda results were encouraging, the desired target is human DNA. Human DNA could be more challenging due to its much larger size and correspondingly fewer binding sites per kb. Most experiments using human DNA involved incubating the complexes with all components at 25°C for 10 minutes prior to raising to 37°C. Human DNA samples were run overnight on the OhmX Analyzer to achieve whole genome coverage.

For testing whether CRISPR/Cas9 could specifically block nicking/tagging with human DNA, a region containing the *FXN* repeat expansion causing FRDA was examined [26]. The interval containing the expansion is 2660 bp and there is a 897 bp interval adjacent to that (Figure 1). Because individuals with FRDA may have many pathogenic expansions in this region that are around 900 bp, as found with GM16798, the adjacent 897 bp interval can sometimes interfere with proper alignment of tags. A gRNA with the sequence GGAAGGACTCTGATTCCACGNGG should uniquely bind to this BssSI recognition sequence and prevent nicking at the site. This is demonstrated in Figure 1 where tags at that site are specifically lost in the blocked sample while alignment occurs at the site in the unaffected GM24694 DNA and the affected GM16798 DNA but not at the blocked GM16798. The blocked sample’s alignment is cleaner than the unblocked sample and the repeat expansion can be clearly seen.

**Figure 1:**
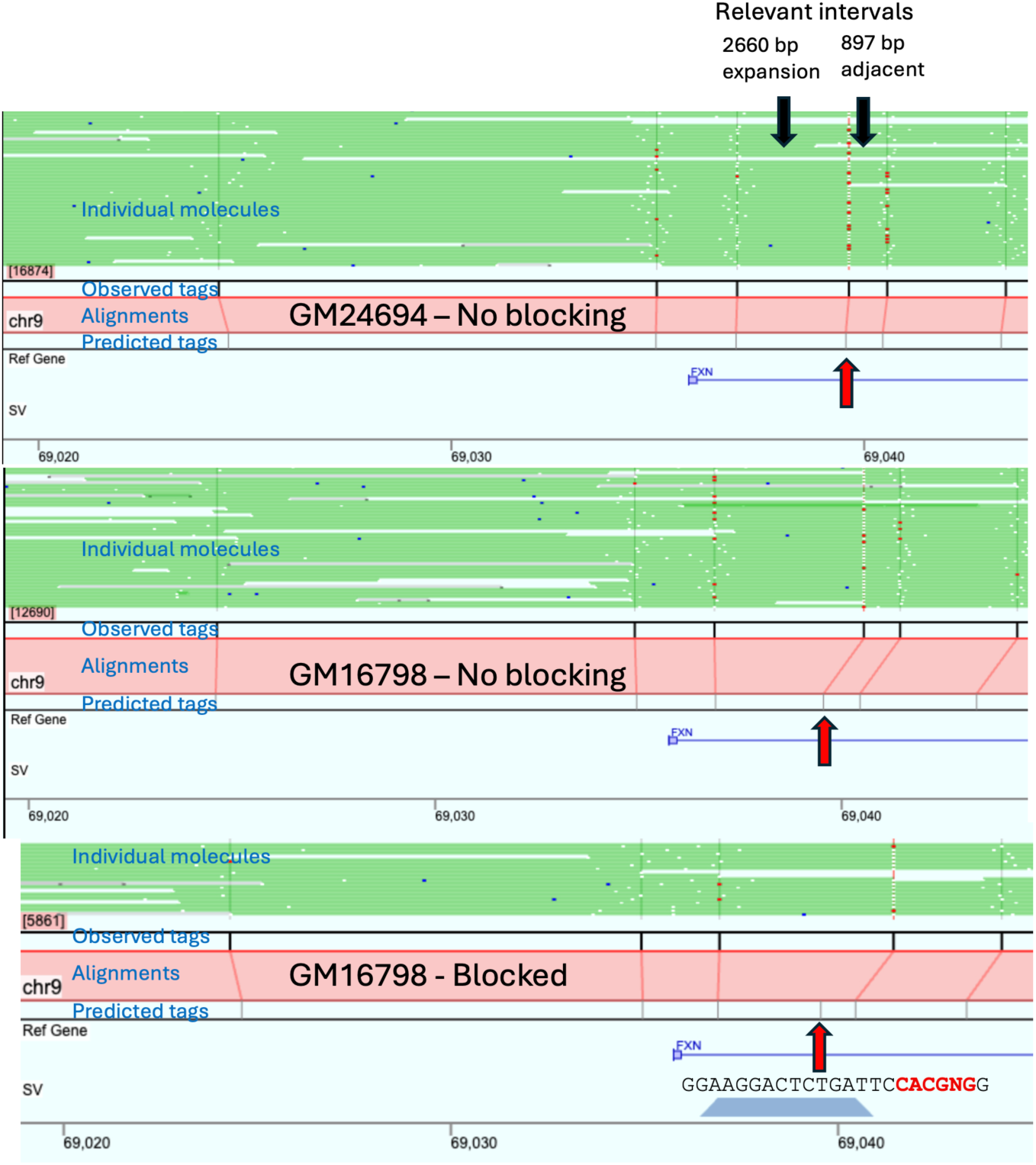
Specific blocking of nick site adjacent to the *FXN* repeat. Individual molecules within the whole-genome assemblies are shown as green lines with each line corresponding to an individual molecule. Tags detected on each molecule are shown by white dots (tag aligned to a called site), red dots (tag missing from a called site), or blue dots (false positive appearing independent of a called site). The genomic coordinates are shown on the bottom of each panel with the predicted tag locations shown above that. The red lines connect the predicted tags below to the observed tags above. If they match, they are connected by a line. The red arrows show the predicted tag sites affected by blocking. DNA is from the Coriell cells (GM24694 and GM16798) indicated in each panel with the bottom panel also including Cas9 and the indicated gRNA.

Typically, choosing a gRNA for specific actions like editing genomic DNA involves selecting sequences that have a single perfect match and as few sites as possible with one or two mismatches [27]. However, our interest was not in looking at a single site but in examining hundreds or thousands of sites simultaneously. Thus, we chose gRNAs using the opposite rationale: gRNAs must have multiple sites with perfect matches and many more sites with mismatches, so that the impact of mismatches on binding could be studied. In addition, we chose sequences that include or overlap a nickase recognition site so that the CRISPR binding effects on the nickase could be more easily monitored. Sequences were also selected to include both isolated and closely paired nick sites which were used for different analyses (Figure 2). The isolated sites were used to study the effect of mismatches and off-target binding. The paired sites were used to study the physical impact of the bound complex on proximal nickase activity. We additionally restricted the analysis to sites where both isolated and paired sites were at least 800 bp away from any other nick site to avoid potential tag misalignments. Many candidate gRNAs met these criteria.

**Figure 2:**
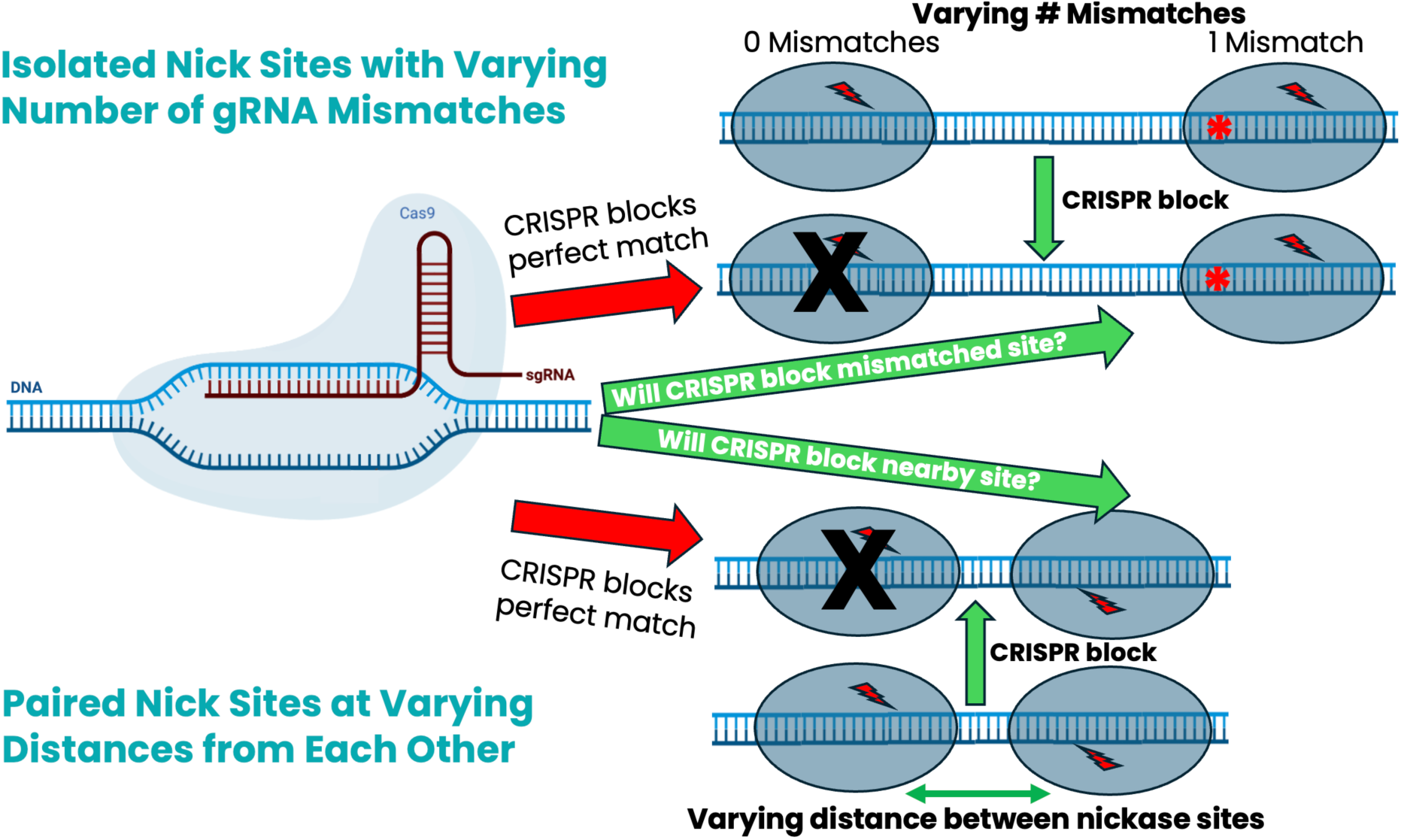
Schematic diagram of dCas9-induced blocking of DNA nickases. dCas9 was used in two modes to examine CRISPR activity. As shown in the top part of the figure, nicking at sites with zero or one mismatch was compared to assess dCas9 binding. In the lower part of the figure, the behavior of pairs of closely spaced nick sites was examined to determine the physical extent to which bound dCas9 can impact nearby activity.

Isolated single-nick sites can be examined in two ways, using either dCas9 or D10A Cas9. The dCas9 approach is useful when a repetitive sequence is being examined, and a nickase recognition site can be placed within or overlapping the gRNA. The disappearance of that nick site can then be monitored. When gRNA sites that do not contain a nickase site are studied, D10A Cas9 can be employed, and the appearance of novel nicks can then be detected. The work presented here uses the dCas9 approach and looks only at the impact of its binding on the nicking of known, pre-existing sites.

To allow examination of thousands of similar sites, we chose to study a repetitive region upstream of the *FMR1* repeat expansion. This region contains a sequence from the THE1C family of transposable elements [28]. The *FMR1* sequence as well as representative similar sequences from throughout the genome are shown in Figure 3. Thousands of other similar sequences are present elsewhere in the genome. The sequences or reverse complements of the gRNAs tested with human DNA are also shown in Figure 3.

**Figure 3:**
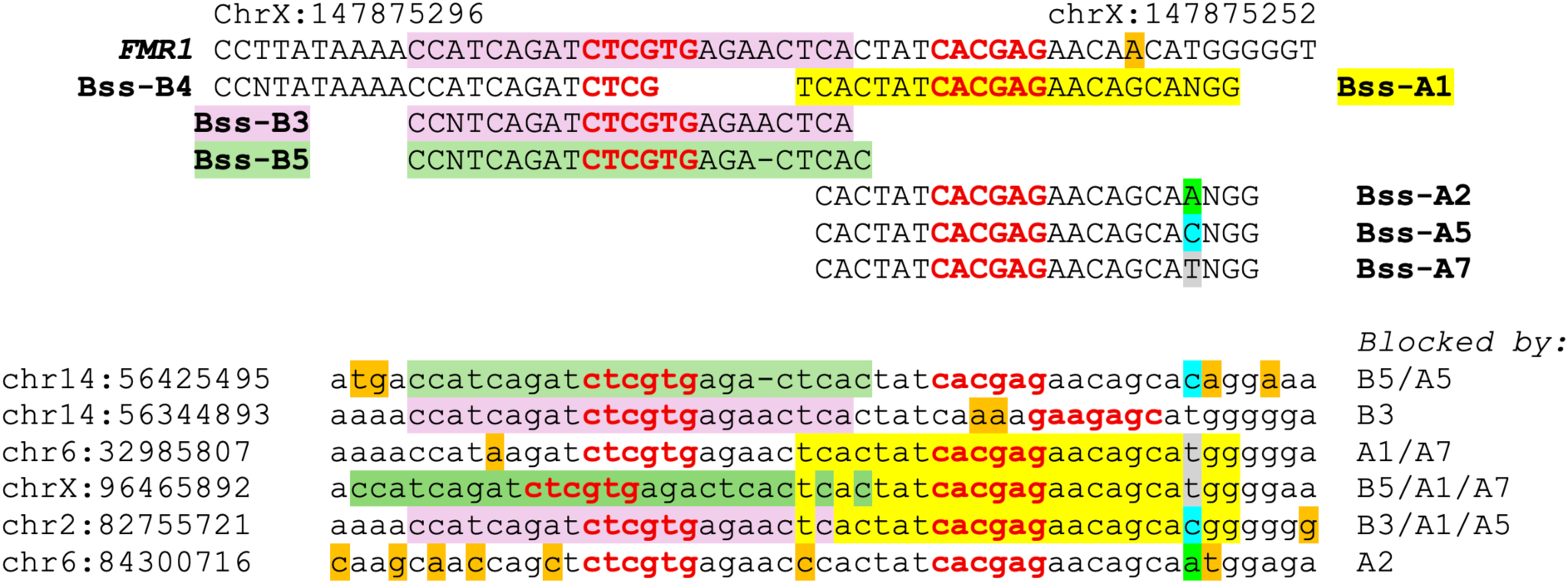
gRNAs upstream of *FMR1* and related sequences. Nt.BspQI and Nb.BssSI recognition sites are in bold red text. gRNA (and the adjacent PAM) and *FMR1* sequences are in capital letters while repetitive segments from other parts of the genome are in lowercase. The gRNA names are in bold black text. The locations of some gRNAs and their binding sites are shaded in different colors. Single bases that do not match the most common sequence are shaded orange. The hg38 genomic start point coordinates for each sequence are shown on the left. gRNAs that should block a given sequence are listed on the right. These sequences represent only a small sample of those similar to the *FMR1* region sequences.

Nearly all perfect genomic matches to these gRNAs bound the CRISPR/Cas9 complex as evidenced by a significant reduction in tagging, going from ∼90-95% to ∼5-10% tagging. Examples of molecules mapped and assembled to the human genome are shown in Figure 4. HG002 DNA was incubated with Cas9 and either Bss-A1 or Bss-A5 and molecular reads processed as normal and assembled using HCE. The assemblies show individual molecules in the region around chr1:76,080,000-76,110,00 on the left and chr11:26,230,000-26,250,000 on the right. The top left tag at 76,097,359 is blocked by Bss-A1 so there are no tags at the designated location. In the panel below, nearly all molecules have a tag (white dots) with just a few missing the tag (red dots). The opposite effect is seen in the right panels where Bss-A1 does not block with a handful of red dots appearing in the top panel while the lower panel is blocked by Bss-A5. Because of the well-characterized gRNA-dependent binding of dCas9, blocking was expected. Fortuitously, the diversity of the related sequences provided an opportunity to examine how mismatches affect binding.

**Figure 4:**
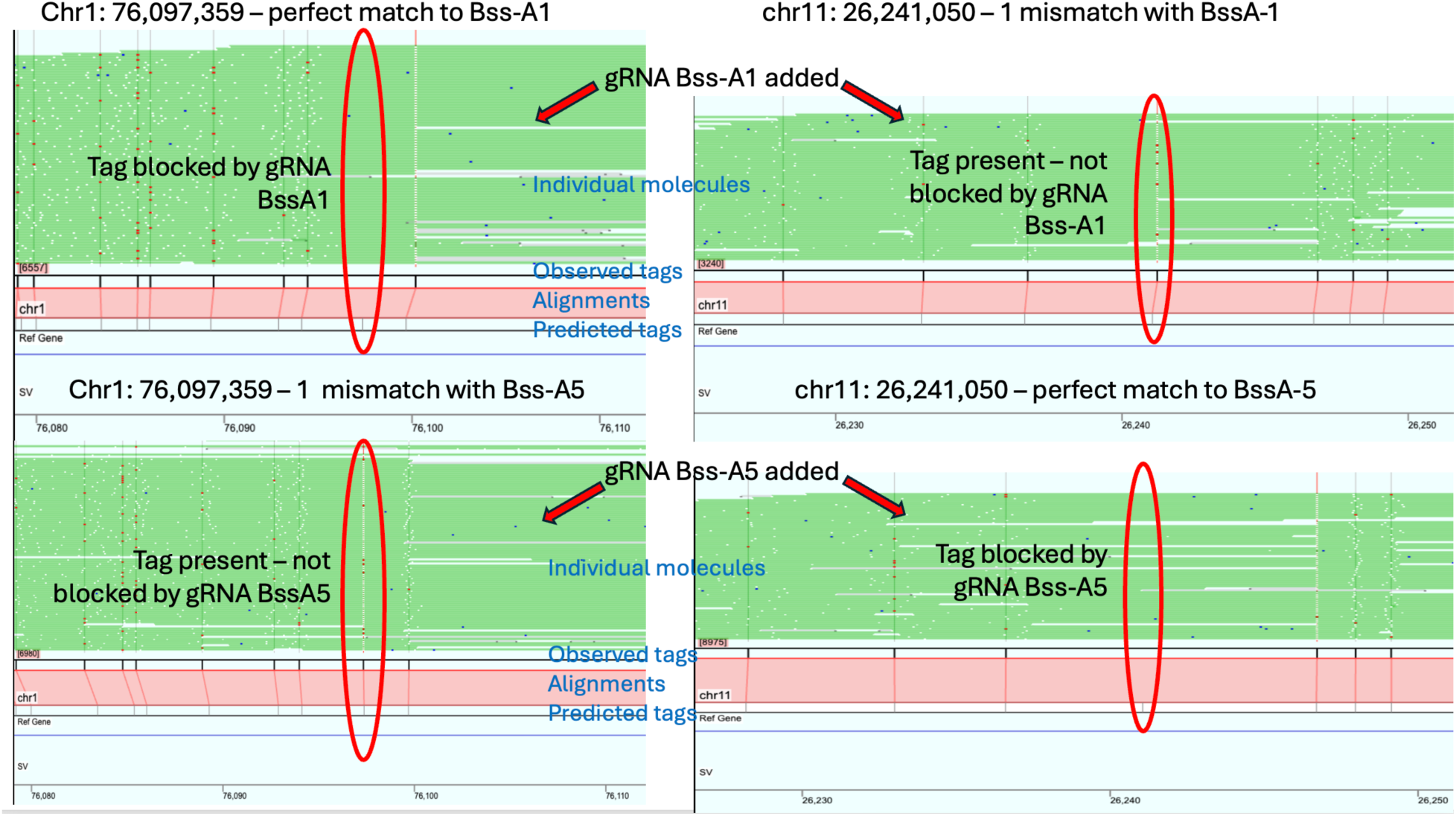
Specific blocking of human DNA with Bss-A1 and Bss-A5. Individual molecules within the whole-genome assemblies are shown as green lines with each line corresponding to an individual molecule. Tags detected on each molecule are shown by white dots (tag aligned to a called site), red dots (tag missing from a called site), or blue dots (false positive appearing independent of a called site). The genomic coordinates are shown on the bottom of each panel with the predicted tag locations shown above that. The red lines connect the predicted tags below to the observed tags above. If they match, they are connected by a line. The top panels are from DNA treated with gRNA Bss-A1 and the bottom with Bss-A5. The relevant potentially blockable tags are enclosed within a red oval.

Because of the danger that misaligning tag sites would have on interpretation, only mismatched gRNA binding sites that were located at least 800 bp away from any other predicted tag site were examined. All possible single-base mismatches that did not affect the overlapping Nb.BssSI recognition sites were evaluated. As shown in Supplementary Table 1, there are examples of most changes somewhere in the genome with the gRNAs examined.

The four gRNAs represented in Supplementary Table 1 and Figure 5 are nearly identical. Positions are numbered with 1 being most distal to the PAM site and 20 being adjacent to the PAM site NGG. Bss-A1 is offset from the other three by a single base so most Bss-A1 binding sites will also bind one of the other three gRNAs. The three offset gRNAs are identical except for position 20 which is in the seed region and corresponds to the PAM N position in Bss-A1. The positions where the BssSI site is located (8-13 for Bss-A1 and 7-12 for the others) could not be examined for mismatches because such mismatched sites would not nick regardless of whether the gRNA was bound or not. Out of the 51 other mismatches that could potentially occur for each gRNA, single-mismatch data was generated for 37, 35, 33, and 38 possibilities in Bss-A1, Bss-A2, Bss-A5, and Bss-A7, respectively. For each position, Supplementary Table 1 reports the median nicking frequency for each alternate base and the combined median across bases; the summary is shown in Figure 5. None of the positions that have at least three sequence examples in the first 15 positions of any of the gRNAs has more than 20% median nicking, indicating that there is significant dCas9 binding at every such mismatch. At positions 16-18, there is more nicking and hence less binding of the CRISPR complex due to these mismatches. Positions 19 and 20 show even higher levels of nicking, indicating a greater importance for stable binding.

**Figure 5:**
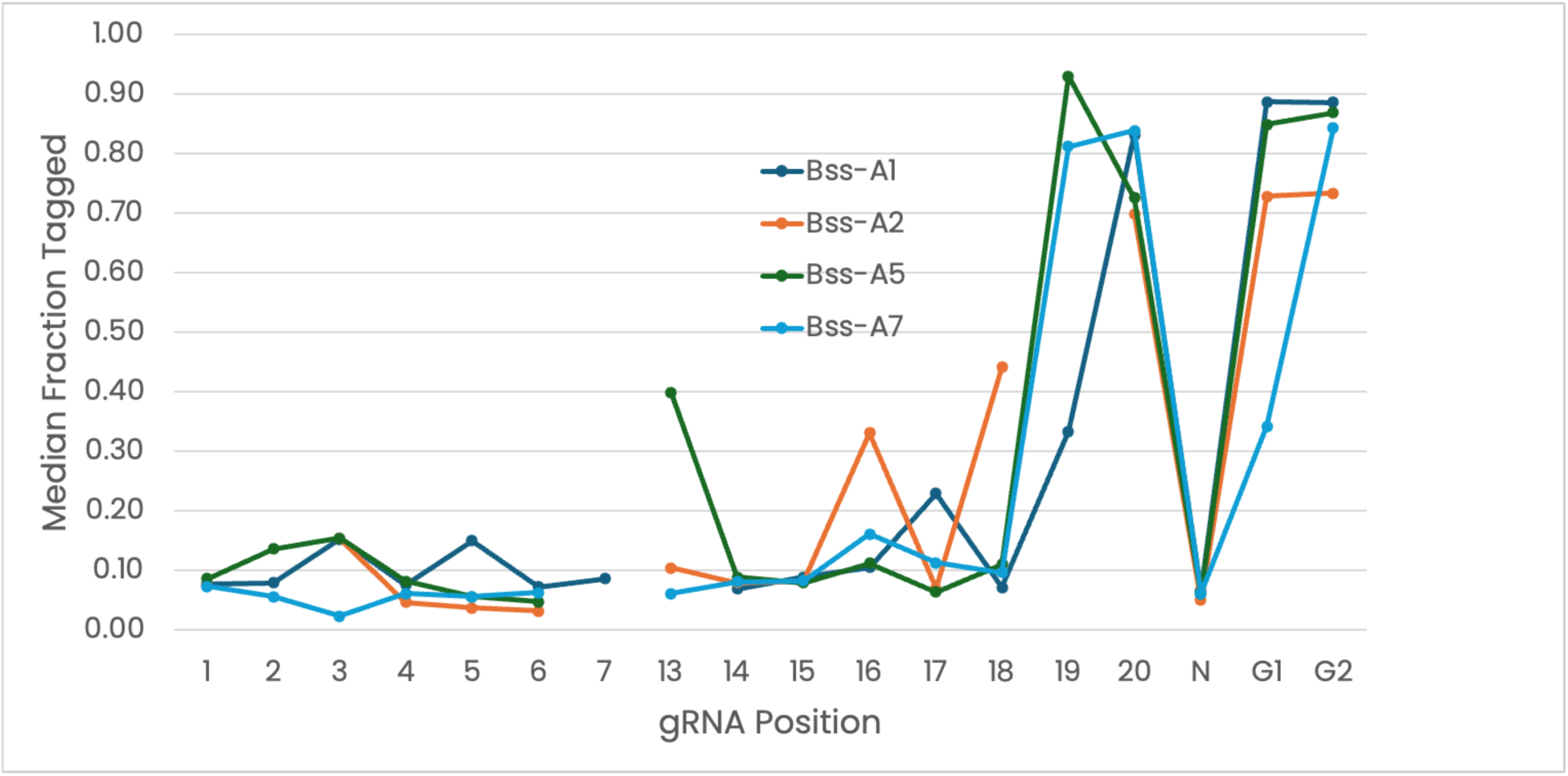
Median tagging frequency for mismatched genomic sequences. Median tagging frequency for all positions with four gRNAs, Bss-A1, Bss-A2, Bss-A5, and Bss-A7, is listed starting with position 1 most distal from the PAM region. Mismatches within the Nb.BssSI recognition site could not be measured (positions 8-13 for Bss-A1 and 7-12 for the other gRNAs). The complete data set is provided in Supplementary Table 1.

The N position in the PAM sequence makes little or no difference to binding. A mismatch in either G in the PAM leads to full nicking, indicating no CRISPR binding, reinforcing the importance of these bases in the process. One possibility for reducing nicking when there is a mutated PAM would be to use an alternate PAM sequence at an adjacent or nearby position. G1 and G2 mismatches were separated into categories based on whether there is a second potential PAM site within 3 bp of the original PAM site. No significant difference in median tagging frequency was observed with or without an alternate NGG (Table 1).

**Table 1:** Median tagging frequency for G1 and G2 mismatched sequences. The regions within 3 bp of all sequences with a mismatch at either the G1 or G2 position in the PAM were inspected for alternate NGG sequences. Median tagging frequencies were determined for those with no alternate NGG and those with alternate NGGs up to 3 bp away in either direction.

| Alternate PAM Distance | # | Median Tagging % |
| --- | --- | --- |
| No other PAM | 23 | 84.2 |
| +1 bp | 17 | 85.7 |
| +2 bp | 13 | 87.1 |
| -3 to +3 bp | 43 | 86.2 |

Isolated nick sites can provide information about binding and activity, whereas pairs of closely spaced nick sites provide a very different type of information. Depending how dCas9 binds the DNA, the paired sites can have three different outcomes. If neither site is bound by dCas9, both positions are nicked. When two such nicks are close together and on opposite strands, the double-nicking can produce a double-stranded cut, causing a Proximity Break (PB) and terminating contig assembly. If only a single site is bound and blocked, the DNA will be intact, and the other site will still have a tag. If both sites are bound and blocked, the DNA will be intact with no tags. Because the repetitive sequences shown in Figure 4 diverge significantly across the human genome, many distances between nick sites can be interrogated. The paired sites were filtered with the same 800 bp criterion used with isolated sites to ensure correct tag alignment.

Among the gRNAs tested, the smallest gap between the end of the gRNA and the nearest non-overlapping nick site was 4 bp. Distances up to 14 bp were examined. As shown in Table 2 and Figure 6, recognition sites 4-7 bp from the gRNA are significantly affected by the nearby bound CRISPR complex, with reduced tagging observed in most molecules 4 – 6 bp from the targeted site. When the second recognition site is separated from the end of the gRNA sequence by 8 bp or more, there is little or no effect on nicking. This is shown graphically in Figure 6.

**Figure 6:**
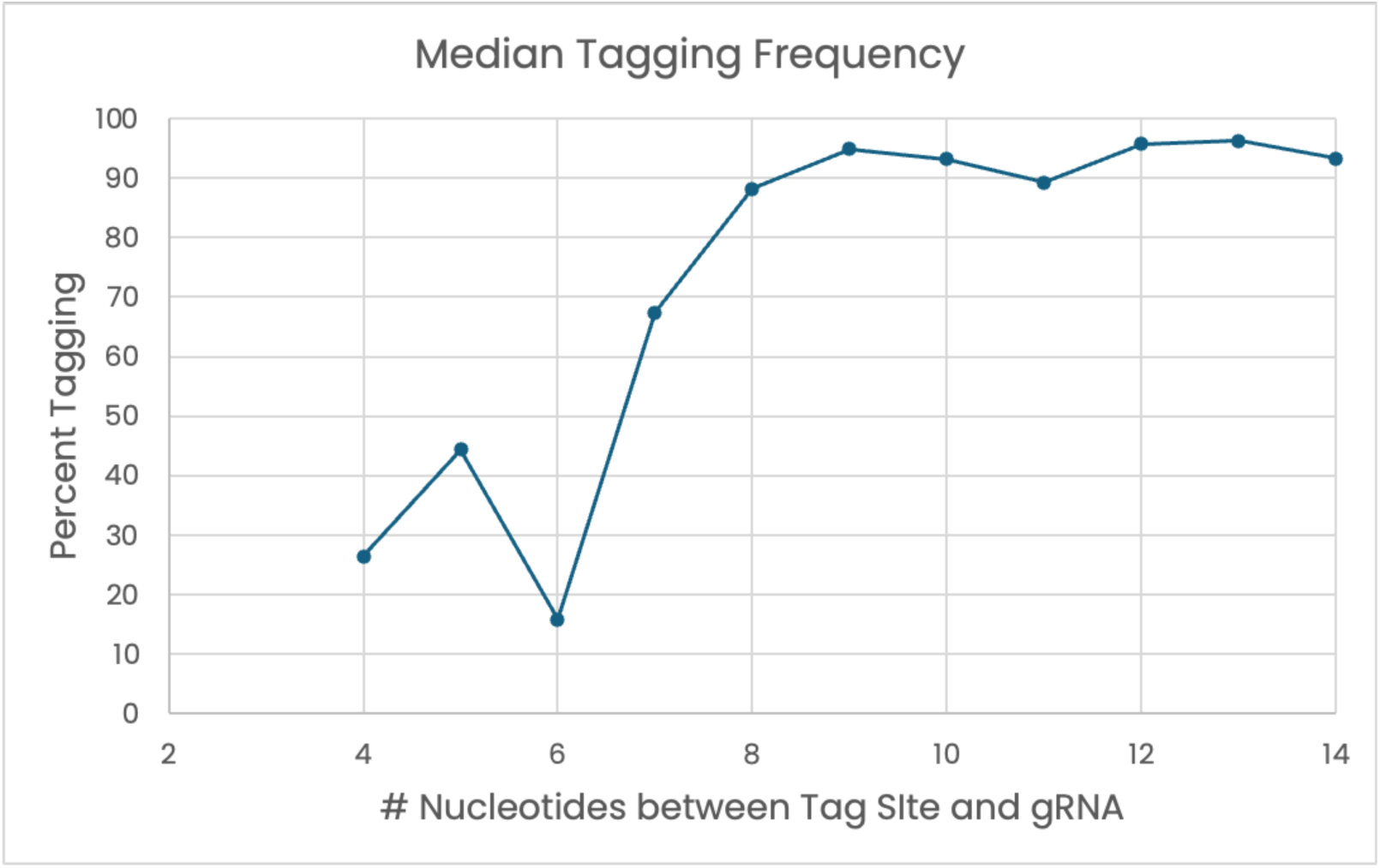
Length dependence of median tagging frequency for paired sites. The combined data for all gRNAs in Table 2 are shown graphically.

**Table 2:** Median tagging frequency as a function of distance from gRNA. The median tagging frequency for all paired sites with a 4-14 bp separation between the nearest nick site and the end of six gRNAs is listed. The number of sites examined for each distance is provided for the individual gRNAs and for all gRNAs combined. The gRNA sequences are shown in Figure 4.

| Distance from gRNA (bp) | BssS-B3 0C Pre-incubation |  | BssS-B3 25C Pre-incubation |  | BssS-B5 |  | BssS-A1 |  | BssS-A2 |  | BssS-A5 |  | BssS-A7 |  | All |  |
| --- | --- | --- | --- | --- | --- | --- | --- | --- | --- | --- | --- | --- | --- | --- | --- | --- |
|  | # | Median % Tagged | # | Median % Tagged | # | Median % Tagged | # | Median % Tagged | # | Median % Tagged | # | Median % Tagged | # | Median % Tagged | # | Median % Tagged |
| 4 |  |  |  |  | 3 | 27.8 | 1 | 18.5 |  |  |  |  |  |  | 4 | 26.4 |
| 5 | 19 | 55.4 | 19 | 41.7 |  |  |  |  |  |  |  |  | 1 | 40.5 | 39 | 44.4 |
| 6 |  |  |  |  |  |  | 14 | 12.4 | 2 | 75.2 | 1 | 21.6 | 3 | 73.5 | 20 | 15.8 |
| 7 |  |  |  |  |  |  |  |  |  |  | 1 | 79.7 | 14 | 67.2 | 15 | 67.4 |
| 8 |  |  |  |  | 8 | 86.9 | 1 | 91.7 |  |  |  |  |  |  | 9 | 88.2 |
| 9 | 46 | 96.8 | 47 | 96.8 |  |  | 42 | 89.6 |  |  |  |  | 1 | 92.1 | 136 | 94.9 |
| 10 | 4 | 94.8 | 4 | 93.2 |  |  | 24 | 95.3 | 5 | 96.1 | 21 | 88.7 | 31 | 92.0 | 89 | 93.2 |
| 11 | 1 | 86.6 | 1 | 100 |  |  |  |  |  |  | 5 | 85.1 | 29 | 90.2 | 36 | 89.2 |
| 12 | 28 | 94.3 | 28 | 96.8 |  |  |  |  |  |  |  |  |  |  | 56 | 95.7 |
| 13 |  |  |  |  | 1 | 96.3 |  |  |  |  |  |  |  |  | 1 | 96.3 |
| 14 | 2 | 91.3 | 2 | 97.5 |  |  |  |  |  |  |  |  |  |  | 4 | 93.3 |

## Discussion

The 2020 Nobel Prize in Chemistry was historic in several aspects. Firstly, it was awarded to Jennifer A Doudna and Emmanuelle Charpentier for their landmark 2012 publication on the programmable dual RNA guided DNA endonuclease for genome editing [5]. Secondly, it was awarded just eight years after the original publication, highlighting CRISPR/Cas9 methodology as revolutionary for global research. Subsequently, the clinical research community followed with multiple pre-clinical research and clinical trials to demonstrate its role in cancer therapeutics (https://clinicaltrials.gov/study/NCT06783270). The FDA approval of Casgevy to treat patients with Sickle cell disease, in 2023, marked another milestone for genome editing community indicating the innovative advancement of the CRISPR mediated approaches for gene therapy [29].

The results presented here demonstrate that EGM can be used to generate detailed information about the activity of DNA-modifying proteins. Even though EGM is a genome mapping technology with resolution on the order of hundreds of base pairs, judicious study design enables functional differences in protein activity to be resolved at base-pair resolution. The ability to simultaneously assess the whole genome provides broader information than available with many other technologies. The flexibility of being able to add or block nick sites nearly at will provides many opportunities to study DNA-modifying proteins and their interactions with DNA and other proteins.

Using different technologies to assess the impact of mismatches on CRISPR/Cas function is complicated by the myriad ways in which the RNP is used. Some methods, such as CRISPR interference, require only the binding activity of the complex, while others, such as direct chemical sequence alteration, require both binding and nuclease activity. Prime editing and CRISPR activation require yet other properties [30]. Even when only knowledge about binding activity is required, outputs from different technologies may assess different aspects of the binding process. For example, massively parallel arrays examine stable binding to oligos[31] and Förster resonance energy transfer (FRET) provides oligo-binding kinetics [32]. Other assays require DNA cleavage to yield a positive result [33]. The EGM assays described here require only binding and not subsequent cleavage of DNA. This binding must be stable as a rapid association/dissociation cycle would allow nickases access to the sites. Furthermore, the CRISPR/Cas interaction with the targeted genomic DNA is more authentic with EGM in that the target is full-length DNA rather than simply oligos. The full diversity of the human genome sequence is present during the binding though the typical cellular complement of DNA binding proteins is not. As a result, it is not surprising that the EGM output is consistent with other technologies, though results may not be identical to those obtained from methods using oligos and a limited target space or carried out *in vivo*.

While we have been using CRISPR as a tool to customize and improve EGM assays, it is also possible to use EGM to better understand CRISPR/Cas systems and improve their functional utility. The distance at which DNA nickases are inhibited by CRISPR/dCas9 binding was resolved at base-pair resolution with 4-7 bp distances showing significant inhibition of tagging due to CRISPR binding. In contrast, distances of 8 bp and greater show minimal impact. This short-range effect is consistent with CRISPR/Cas9 structural studies that show a DNA-protected region of about 30 bp [34]. Extrapolating this distance information allows a more informed design of other gRNAs that should be effective across a variety of sequence contexts. It is also noteworthy that the strongest impact on binding strength for this protein-dependent annealing is at one end of the RNA. This contrasts with earlier work on a comparable A-form Locked nucleic acid (LNA)/DNA duplex in the absence of protein, where the positions most important for stability were in the middle of the duplex [35].

The ability of CRISPR complexes to bind DNA at lower temperatures than many enzymes provides flexibility, allowing components to be added simultaneously to simplify protocols. When CRISPR is used to block nickases or other enzymes that require higher temperatures, it can be added at the same time as the enzymes being tested because those enzymes may not be active at low temperatures. CRISPR binding occurs before nicking can take place. The temperature can then be shifted without additional mixing steps. This allows the separation of binding activity from functional activity.

In this work, we have focused on blocking pre-existing nick sites with dCas9, but it is also possible to create new tagging sites via D10A Cas9. This opens up a wider range of gRNA sites that can be examined for the appearance of new tags at on- and off-target sites. We have focused on a single Cas protein, but these methods are easily extended to other Cas homologs and isoforms in which different proteins could be examined under the same conditions or at altered times and temperatures. The method offers the advantage of examining the whole genome experimentally, rather than relying on predictive models, which may overlook unappreciated factors. Long range interactions are also taken into account with the EGM approach though the impact of cellular DNA binding proteins cannot be observed.

The information derived from this work may help the design of fusions with the active domains of other proteins. Such fusion proteins will further enhance the range of activities that can be envisioned with the DNA-bound complexes [9]. Thus, understanding how CRISPR interacts or blocks the activity of other proteins is important for predicting how well combinations of proteins or domains may work together.

Although this original work is purely based on technology evaluation and a feasibility study, we also demonstrated the potential of this methodology for its impact in repeat expansion diseases, such as FRDA. Our Coriell cell line data demonstrates significant promise and potential. Additional studies from clinical research subjects are warranted to evaluate the CRISPR mediated EGM approach for its utility in the accurate sizing of the FXN repeats that can discern the normal range of repeats (5-33 GAA repeats) from the intermediate (34-65 GAA repeats) and pathogenic (66-1300 GAA repeats) alleles.

## Materials and Methods

Single-guide RNAs (sgRNAs) and Cas9 proteins were purchased from Integrated DNA Technologies (IDT). The two components were combined in equimolar amounts and mixed with PBS to create ribonucleoprotein (RNP) complexes after incubating at room temperature for 10 minutes as recommended by IDT. Lambda DNA and enzymatic nickases (Nt.BspQI and Nb.BssSI) were purchased from New England Biolabs (NEB). With all reactions, albumin-free buffers were used. Except where noted, human DNA was from HG002 cells. All human DNA was extracted from cells using the NEB Monarch HMW DNA Extraction Kit for Cells & Blood with the recommended protocol modified to use an agitation speed of 1200 rpm during cell lysis. Cell lines were obtained from Coriell, grown as recommended, frozen in 1-million-cell aliquots, and stored at -80°C prior to use.

The EGM and genome preparation methods for the OhmX Platform are described at https://www.nabsys.com/learn/sample-prep-kit. Briefly, the standard protocol starts with nicking ultra-long DNA and adding biotinylated dNTPs to the nick sites during the Simultaneous Nicking and Labeling step. Next, the labeled DNA is purified to remove unincorporated dNTPs. This DNA is then tagged with a streptavidin-DNA complex, followed by coating the tagged DNA with RecA protein to straighten and stiffen it. This complex is injected into the OhmX Analyzer and data collected over minutes to hours, depending on the coverage needs. Data is collected as voltage versus time and then converted into physical distances for use in mapping to a reference or assembling into contigs using High-Definition Mapping (HDM) Analysis software and Human Chromosome Explorer (HCE) software, respectively.

When testing CRISPR/Cas9 systems for blocking nick sites, dCas9 RNP complexes are combined with all standard reagents for the Simultaneous Nicking and Labeling protocol. The RNPs were tested under a variety of incubation times and temperatures and under different reaction assembly conditions, including performing the RNP incubation step before adding the nicking enzymes. Ultimately, the dCas9 blocking reaction was performed by adding the RNP complexes directly to the Simultaneous Nicking and Labeling reaction and incubating at 25°C for 10 minutes before moving the reaction to 37°C for the 1-hour nicking and labeling step. All other steps of the EGM sample prep protocol were followed according to the standard conditions.

## Supporting information

Supplemental mismatch data

Supplemental length data

## Acknowledgements

We thank the many Nabsys employees who have developed the assays and analytics that allowed us to carry out this work.

## Author contributions

LSO designed, developed, and carried out the CRISPR assays for use with EGM, helped with analyzing data, and writing the manuscript, WH developed and carried out the mismatch assays, and helped analyze data and write the manuscript, SK optimized CRISPR assays and helped write the manuscript, KLM helped write and revise the manuscript, and JFT helped design the assays, analyze the data, and write the manuscript.

## Data availability

Data is supplied in supplementary tables.

## Supplementary Information

**Supplementary Figure 1:**
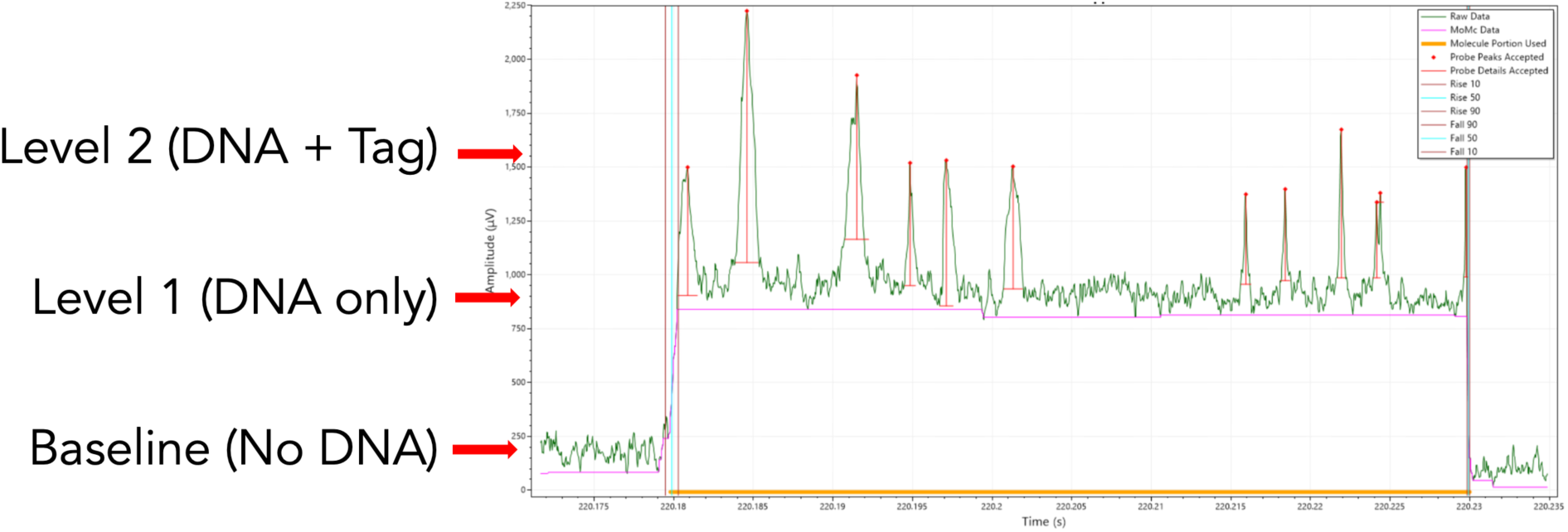
Voltage patterns for lambda DNA. The voltage traces for single molecules of tagged lambda molecules are shown as a function of time. Either end of a molecule may enter the channel first. When the DNA transits the detector region, the voltage changes from baseline due to the current blockade (Level 1). It changes again whenever a tag is present (Level 2). Molecules speed up as they traverse the nanochannel. The time between tags is measured and converted to base pairs using a signal processing algorithm that accounts for differential speed as the DNA accelerates through the nanochannel.

**Supplementary Table 1:**
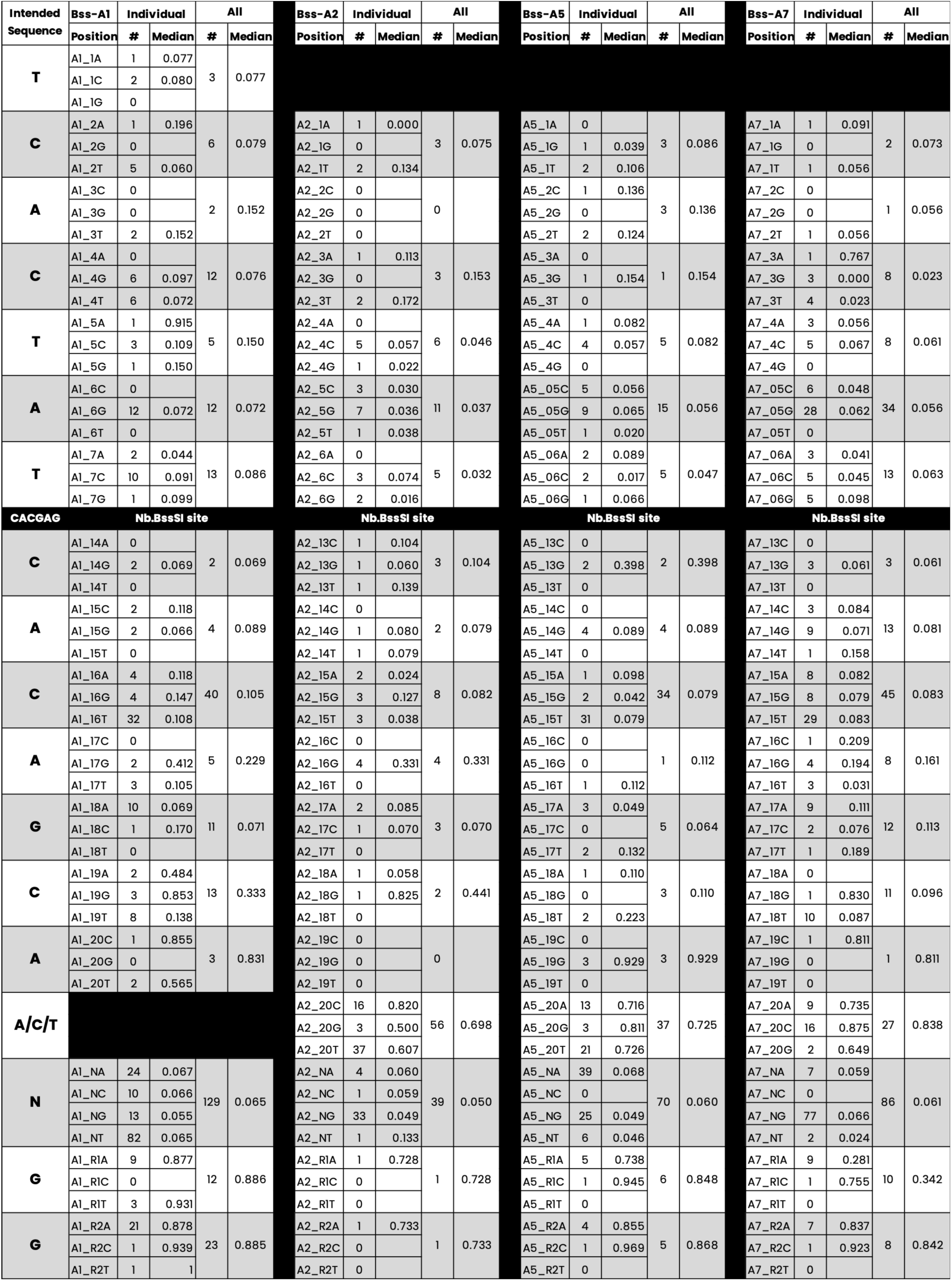
Median tagging frequency for mismatched genomic sequences. Median tagging frequency for all positions with four gRNAs, Bss-A1, Bss-A2, Bss-A5, and Bss-A7, is listed starting with position 1 most distal from the PAM region. Bss-A1 is offset by one base relative to the other three so its data is offset by one in the table. The number of examples for each substitution is shown in the second column and the median frequency in the third. Each set of mismatched bases is combined in the next column.

