## Supplemental mismatch data for "CRISPR/Cas9 Optimization Using Electronic Genome Mapping: Potential Role for Studying Human Genetic Disease"

| Mismatch | Chromosor | hg38 Position | Tags | Molecules | % Tagged |
|---|---|---|---|---|---|
| A1_No_Mismatch | chr1 | 3,575,739 | 11 | 69 | 15.9 |
| A1_6G | chr1 | 4,099,321 | 2 | 56 | 3.6 |
| A1_R2A | chr1 | 4,669,530 | 91 | 100 | 91.0 |
| A1_No_Mismatch | chr1 | 6,976,874 | 3 | 41 | 7.3 |
| A1_No_Mismatch | chr1 | 7,196,126 | 3 | 41 | 7.3 |
| A1_R1T | chr1 | 9,020,840 | 67 | 72 | 93.1 |
| A1_No_Mismatch | chr1 | 11,113,896 | 2 | 34 | 5.9 |
| A1_4T | chr1 | 13,847,587 | 5 | 53 | 9.4 |
| A1_19T | chr1 | 14,428,766 | 8 | 68 | 11.8 |
| A1_No_Mismatch | chr1 | 17,704,400 | 3 | 46 | 6.5 |
| A1_No_Mismatch | chr1 | 20,132,536 | 7 | 56 | 12.5 |
| A1_No_Mismatch | chr1 | 25,333,875 | 1 | 45 | 2.2 |
| A1_18A | chr1 | 27,805,052 | 2 | 55 | 3.6 |
| A1_19G | chr1 | 32,504,995 | 59 | 77 | 76.6 |
| A1_2T | chr1 | 33,239,677 | 11 | 100 | 11.0 |
| A1_No_Mismatch | chr1 | 33,547,977 | 2 | 80 | 2.5 |
| A1_7C | chr1 | 34,671,038 | 10 | 99 | 10.1 |
| A1_No_Mismatch | chr1 | 40,505,889 | 5 | 77 | 6.5 |
| A1_No_Mismatch | chr1 | 47,596,180 | 4 | 79 | 5.1 |
| A1_No_Mismatch | chr1 | 53,341,254 | 4 | 82 | 4.9 |
| A1_7C | chr1 | 54,934,336 | 4 | 73 | 5.5 |
| A1_20T | chr1 | 55,346,206 | 12 | 40 | 30.0 |
| A1_1A | chr1 | 55,621,251 | 6 | 78 | 7.7 |
| A1_19G | chr1 | 56,588,304 | 64 | 69 | 92.8 |
| A1_No_Mismatch | chr1 | 57,938,448 | 8 | 97 | 8.2 |
| A1_4G | chr1 | 58,513,155 | 4 | 37 | 10.8 |
| A1_No_Mismatch | chr1 | 59,031,497 | 4 | 103 | 3.9 |
| A1_7C | chr1 | 59,378,273 | 13 | 76 | 17.1 |
| A1_16T | chr1 | 67,509,309 | 11 | 40 | 27.5 |
| A1_16A | chr1 | 71,854,787 | 6 | 38 | 15.8 |
| A1_No_Mismatch | chr1 | 76,097,359 | 2 | 73 | 2.7 |
| A1_20C | chr1 | 78,122,601 | 53 | 62 | 85.5 |
| A1_16A | chr1 | 79,251,486 | 2 | 43 | 4.7 |
| A1_R2T | chr1 | 81,714,076 | 83 | 83 | 100.0 |
| A1_16T | chr1 | 82,610,368 | 1 | 32 | 3.1 |
| A1_16T | chr1 | 83,586,976 | 2 | 41 | 4.9 |
| A1_No_Mismatch | chr1 | 87,503,056 | 2 | 74 | 2.7 |
| A1_R2A | chr1 | 90,607,425 | 73 | 76 | 96.1 |
| A1_No_Mismatch | chr1 | 90,757,650 | 5 | 68 | 7.4 |
| A1_R1A | chr1 | 90,862,991 | 64 | 68 | 94.1 |

| | | | | | |
|---|---|---|---|---|---|
| A1_No_Mismatch | chr1 | 94,428,597 | 2 | 53 | 3.8 |
| A1_R2A | chr1 | 95,629,434 | 56 | 62 | 90.3 |
| A1_19T | chr1 | 96,881,349 | 14 | 38 | 36.8 |
| A1_1C | chr1 | 101,979,428 | 4 | 82 | 4.9 |
| A1_18A | chr1 | 112,150,403 | 2 | 32 | 6.3 |
| A1_No_Mismatch | chr1 | 114,194,161 | 7 | 110 | 6.4 |
| A1_No_Mismatch | chr1 | 117,983,253 | 1 | 29 | 3.4 |
| A1_5A | chr1 | 146,694,255 | 86 | 94 | 91.5 |
| A1_7C | chr1 | 151,868,313 | 17 | 85 | 20.0 |
| A1_No_Mismatch | chr1 | 157,674,503 | 6 | 53 | 11.3 |
| A1_No_Mismatch | chr1 | 157,759,091 | 1 | 73 | 1.4 |
| A1_16T | chr1 | 159,995,542 | 8 | 78 | 10.3 |
| A1_No_Mismatch | chr1 | 161,754,705 | 8 | 93 | 8.6 |
| A1_No_Mismatch | chr1 | 167,597,384 | 7 | 72 | 9.7 |
| A1_No_Mismatch | chr1 | 167,884,402 | 6 | 48 | 12.5 |
| A1_No_Mismatch | chr1 | 168,464,451 | 2 | 44 | 4.5 |
| A1_R2A | chr1 | 169,757,664 | 55 | 60 | 91.7 |
| A1_R1A | chr1 | 170,834,490 | 33 | 34 | 97.1 |
| A1_R1A | chr1 | 171,126,234 | 30 | 31 | 96.8 |
| A1_R2C | chr1 | 171,837,868 | 62 | 66 | 93.9 |
| A1_R2A | chr1 | 171,955,956 | 43 | 49 | 87.8 |
| A1_19T | chr1 | 173,340,974 | 8 | 90 | 8.9 |
| A1_R2A | chr1 | 178,024,406 | 54 | 60 | 90.0 |
| A1_5C | chr1 | 179,146,659 | 10 | 92 | 10.9 |
| A1_No_Mismatch | chr1 | 179,302,403 | 47 | 53 | 88.7 |
| A1_R2A | chr1 | 187,507,309 | 27 | 30 | 90.0 |
| A1_No_Mismatch | chr1 | 187,736,148 | 2 | 45 | 4.4 |
| A1_16T | chr1 | 194,265,066 | 6 | 54 | 11.1 |
| A1_16T | chr1 | 197,995,829 | 12 | 55 | 21.8 |
| A1_No_Mismatch | chr1 | 198,355,520 | 4 | 98 | 4.1 |
| A1_No_Mismatch | chr1 | 198,798,136 | 6 | 68 | 8.8 |
| A1_2T | chr1 | 199,678,209 | 6 | 61 | 9.8 |
| A1_2A | chr1 | 218,448,629 | 11 | 56 | 19.6 |
| A1_18A | chr1 | 223,046,719 | 5 | 46 | 10.9 |
| A1_No_Mismatch | chr1 | 230,360,526 | 4 | 73 | 5.5 |
| A1_6G | chr1 | 230,420,545 | 5 | 75 | 6.7 |
| A1_No_Mismatch | chr1 | 232,252,747 | 15 | 66 | 22.7 |
| A1_R2A | chr1 | 233,889,311 | 39 | 51 | 76.5 |
| A1_7C | chr1 | 237,616,936 | 5 | 52 | 9.6 |
| A1_No_Mismatch | chr1 | 238,747,294 | 4 | 16 | 25.0 |
| A1_No_Mismatch | chr1 | 239,507,312 | 7 | 81 | 8.6 |

| | | | | | |
|---|---|---|---|---|---|
| A1_No_Mismatch | chr1 | 240,751,948 | 6 | 66 | 9.1 |
| A1_No_Mismatch | chr1 | 241,537,815 | 37 | 46 | 80.4 |
| A1_No_Mismatch | chr1 | 244,309,325 | 7 | 48 | 14.6 |
| A1_15G | chr10 | 7,418,678 | 2 | 42 | 4.8 |
| A1_6G | chr10 | 12,817,099 | 3 | 52 | 5.8 |
| A1_1C | chr10 | 17,526,198 | 6 | 54 | 11.1 |
| A1_16T | chr10 | 17,786,882 | 8 | 85 | 9.4 |
| A1_7C | chr10 | 19,231,547 | 6 | 70 | 8.6 |
| A1_No_Mismatch | chr10 | 21,246,976 | 5 | 97 | 5.2 |
| A1_6G | chr10 | 22,032,394 | 2 | 56 | 3.6 |
| A1_16T | chr10 | 23,489,230 | 3 | 37 | 8.1 |
| A1_No_Mismatch | chr10 | 29,673,692 | 2 | 106 | 1.9 |
| A1_No_Mismatch | chr10 | 30,756,830 | 10 | 66 | 15.2 |
| A1_No_Mismatch | chr10 | 30,795,791 | 6 | 54 | 11.1 |
| A1_No_Mismatch | chr10 | 32,222,873 | 6 | 63 | 9.5 |
| A1_No_Mismatch | chr10 | 33,359,695 | 3 | 70 | 4.3 |
| A1_16A | chr10 | 33,599,549 | 5 | 62 | 8.1 |
| A1_No_Mismatch | chr10 | 52,674,364 | 6 | 58 | 10.3 |
| A1_No_Mismatch | chr10 | 56,065,057 | 3 | 60 | 5.0 |
| A1_19T | chr10 | 56,210,541 | 18 | 30 | 60.0 |
| A1_19G | chr10 | 58,173,556 | 29 | 34 | 85.3 |
| A1_No_Mismatch | chr10 | 59,053,781 | 2 | 44 | 4.5 |
| A1_15G | chr10 | 59,278,144 | 5 | 59 | 8.5 |
| A1_4G | chr10 | 60,132,370 | 3 | 46 | 6.5 |
| A1_No_Mismatch | chr10 | 61,126,799 | 1 | 34 | 2.9 |
| A1_No_Mismatch | chr10 | 61,191,736 | 0 | 6 | 0.0 |
| A1_No_Mismatch | chr10 | 66,116,907 | 3 | 89 | 3.4 |
| A1_6G | chr10 | 75,376,863 | 1 | 63 | 1.6 |
| A1_16G | chr10 | 76,110,620 | 16 | 74 | 21.6 |
| A1_16T | chr10 | 78,242,163 | 9 | 104 | 8.7 |
| A1_No_Mismatch | chr10 | 82,339,504 | 9 | 43 | 20.9 |
| A1_No_Mismatch | chr10 | 97,167,772 | 5 | 62 | 8.1 |
| A1_No_Mismatch | chr10 | 107,825,297 | 13 | 109 | 11.9 |
| A1_18A | chr10 | 108,185,356 | 9 | 66 | 13.6 |
| A1_19A | chr10 | 109,596,330 | 32 | 46 | 69.6 |
| A1_19T | chr10 | 114,094,082 | 6 | 78 | 7.7 |
| A1_7C | chr10 | 114,510,008 | 5 | 70 | 7.1 |
| A1_16T | chr10 | 115,106,617 | 10 | 77 | 13.0 |
| A1_16T | chr10 | 118,855,898 | 13 | 78 | 16.7 |
| A1_2T | chr10 | 119,232,592 | 2 | 53 | 3.8 |
| A1_No_Mismatch | chr10 | 124,369,574 | 2 | 73 | 2.7 |

| | | | | | |
|---|---|---|---|---|---|
| A1_No_Mismatch | chr10 | 125,253,848 | 5 | 40 | 12.5 |
| A1_5C | chr10 | 128,920,975 | 11 | 63 | 17.5 |
| A1_5C | chr10 | 129,072,162 | 4 | 66 | 6.1 |
| A1_No_Mismatch | chr10 | 130,075,711 | 8 | 37 | 21.6 |
| A1_16T | chr10 | 132,600,850 | 11 | 84 | 13.1 |
| A1_No_Mismatch | chr11 | 2,324,556 | 23 | 68 | 33.8 |
| A1_No_Mismatch | chr11 | 9,190,417 | 3 | 65 | 4.6 |
| A1_3T | chr11 | 12,022,287 | 7 | 29 | 24.1 |
| A1_No_Mismatch | chr11 | 15,499,746 | 5 | 58 | 8.6 |
| A1_No_Mismatch | chr11 | 20,987,611 | 2 | 63 | 3.2 |
| A1_16G | chr11 | 22,930,964 | 4 | 51 | 7.8 |
| A1_No_Mismatch | chr11 | 23,384,179 | 1 | 22 | 4.5 |
| A1_No_Mismatch | chr11 | 26,205,128 | 2 | 41 | 4.9 |
| A1_No_Mismatch | chr11 | 29,441,741 | 6 | 39 | 15.4 |
| A1_7C | chr11 | 29,453,377 | 2 | 36 | 5.6 |
| A1_15C | chr11 | 31,937,696 | 12 | 83 | 14.5 |
| A1_16T | chr11 | 36,338,931 | 11 | 65 | 16.9 |
| A1_No_Mismatch | chr11 | 38,748,213 | 3 | 51 | 5.9 |
| A1_16T | chr11 | 40,078,300 | 4 | 38 | 10.5 |
| A1_No_Mismatch | chr11 | 40,625,502 | 6 | 22 | 27.3 |
| A1_16T | chr11 | 43,477,608 | 9 | 67 | 13.4 |
| A1_No_Mismatch | chr11 | 46,187,047 | 0 | 20 | 0.0 |
| A1_No_Mismatch | chr11 | 48,506,989 | 7 | 38 | 18.4 |
| A1_16T | chr11 | 50,141,889 | 3 | 55 | 5.5 |
| A1_16T | chr11 | 50,336,627 | 5 | 35 | 14.3 |
| A1_16T | chr11 | 57,168,860 | 9 | 60 | 15.0 |
| A1_17T | chr11 | 57,193,356 | 6 | 65 | 9.2 |
| A1_6G | chr11 | 61,153,219 | 3 | 58 | 5.2 |
| A1_No_Mismatch | chr11 | 70,899,240 | 2 | 59 | 3.4 |
| A1_No_Mismatch | chr11 | 74,826,389 | 5 | 52 | 9.6 |
| A1_No_Mismatch | chr11 | 77,853,493 | 3 | 79 | 3.8 |
| A1_16A | chr11 | 81,840,067 | 11 | 71 | 15.5 |
| A1_No_Mismatch | chr11 | 87,829,588 | 11 | 68 | 16.2 |
| A1_No_Mismatch | chr11 | 91,367,596 | 10 | 63 | 15.9 |
| A1_18A | chr11 | 92,494,176 | 3 | 45 | 6.7 |
| A1_No_Mismatch | chr11 | 92,723,820 | 3 | 44 | 6.8 |
| A1_16T | chr11 | 102,126,424 | 6 | 42 | 14.3 |
| A1_4T | chr11 | 104,002,354 | 1 | 18 | 5.6 |
| A1_19A | chr11 | 108,602,978 | 28 | 103 | 27.2 |
| A1_7G | chr11 | 111,118,260 | 7 | 71 | 9.9 |
| A1_14G | chr11 | 111,402,111 | 3 | 60 | 5.0 |

| | | | | | |
|---|---|---|---|---|---|
| A1_No_Mismatch | chr11 | 115,026,553 | 5 | 49 | 10.2 |
| A1_No_Mismatch | chr11 | 116,390,322 | 7 | 90 | 7.8 |
| A1_19T | chr11 | 117,397,731 | 4 | 73 | 5.5 |
| A1_4G | chr11 | 123,667,639 | 4 | 66 | 6.1 |
| A1_No_Mismatch | chr11 | 127,322,008 | 10 | 80 | 12.5 |
| A1_R2A | chr11 | 132,461,875 | 81 | 93 | 87.1 |
| A1_16T | chr11 | 132,508,624 | 5 | 79 | 6.3 |
| A1_7C | chr12 | 8,166,434 | 1 | 55 | 1.8 |
| A1_18A | chr12 | 8,345,537 | 4 | 81 | 4.9 |
| A1_No_Mismatch | chr12 | 13,268,098 | 10 | 75 | 13.3 |
| A1_No_Mismatch | chr12 | 20,286,072 | 5 | 43 | 11.6 |
| A1_18A | chr12 | 21,883,332 | 0 | 48 | 0.0 |
| A1_5G | chr12 | 22,182,619 | 6 | 40 | 15.0 |
| A1_16T | chr12 | 23,746,875 | 4 | 53 | 7.5 |
| A1_No_Mismatch | chr12 | 26,910,465 | 3 | 55 | 5.5 |
| A1_No_Mismatch | chr12 | 28,173,251 | 3 | 32 | 9.4 |
| A1_No_Mismatch | chr12 | 29,867,947 | 4 | 94 | 4.3 |
| A1_No_Mismatch | chr12 | 38,673,073 | 1 | 24 | 4.2 |
| A1_No_Mismatch | chr12 | 39,995,764 | 1 | 39 | 2.6 |
| A1_19T | chr12 | 40,581,079 | 11 | 33 | 33.3 |
| A1_No_Mismatch | chr12 | 41,559,821 | 53 | 54 | 98.1 |
| A1_18A | chr12 | 44,484,721 | 5 | 70 | 7.1 |
| A1_6G | chr12 | 47,401,347 | 5 | 60 | 8.3 |
| A1_No_Mismatch | chr12 | 58,355,080 | 4 | 61 | 6.6 |
| A1_No_Mismatch | chr12 | 58,859,974 | 8 | 86 | 9.3 |
| A1_16T | chr12 | 65,920,588 | 3 | 47 | 6.4 |
| A1_16T | chr12 | 66,511,118 | 3 | 37 | 8.1 |
| A1_18A | chr12 | 72,229,366 | 5 | 51 | 9.8 |
| A1_6G | chr12 | 79,059,171 | 5 | 12 | 41.7 |
| A1_17T | chr12 | 82,620,918 | 4 | 38 | 10.5 |
| A1_16T | chr12 | 87,182,061 | 5 | 32 | 15.6 |
| A1_No_Mismatch | chr12 | 88,019,204 | 6 | 47 | 12.8 |
| A1_17G | chr12 | 89,514,163 | 11 | 42 | 26.2 |
| A1_No_Mismatch | chr12 | 93,045,787 | 7 | 65 | 10.8 |
| A1_No_Mismatch | chr12 | 95,808,268 | 8 | 64 | 12.5 |
| A1_No_Mismatch | chr12 | 98,920,597 | 6 | 81 | 7.4 |
| A1_R2A | chr12 | 99,326,074 | 54 | 62 | 87.1 |
| A1_6G | chr12 | 101,584,775 | 11 | 55 | 20.0 |
| A1_No_Mismatch | chr12 | 101,856,152 | 6 | 61 | 9.8 |
| A1_20T | chr12 | 102,369,029 | 59 | 71 | 83.1 |
| A1_No_Mismatch | chr12 | 102,676,000 | 2 | 29 | 6.9 |

| | | | | | |
|---|---|---|---|---|---|
| A1_16T | chr12 | 102,742,776 | 4 | 64 | 6.3 |
| A1_No_Mismatch | chr12 | 103,555,395 | 3 | 83 | 3.6 |
| A1_16T | chr12 | 104,396,083 | 4 | 72 | 5.6 |
| A1_No_Mismatch | chr12 | 105,927,626 | 4 | 47 | 8.5 |
| A1_No_Mismatch | chr12 | 107,591,524 | 3 | 58 | 5.2 |
| A1_16T | chr12 | 109,639,248 | 3 | 81 | 3.7 |
| A1_3T | chr12 | 110,798,466 | 5 | 78 | 6.4 |
| A1_6G | chr12 | 113,027,846 | 6 | 78 | 7.7 |
| A1_No_Mismatch | chr12 | 115,187,840 | 6 | 64 | 9.4 |
| A1_16T | chr12 | 119,934,898 | 7 | 126 | 5.6 |
| A1_4T | chr12 | 122,160,863 | 4 | 104 | 3.8 |
| A1_4G | chr12 | 123,844,204 | 9 | 59 | 15.3 |
| A1_No_Mismatch | chr12 | 129,744,959 | 5 | 64 | 7.8 |
| A1_17G | chr12 | 131,436,431 | 40 | 71 | 56.3 |
| A1_15C | chr13 | 25,223,725 | 6 | 65 | 9.2 |
| A1_No_Mismatch | chr13 | 35,963,706 | 3 | 91 | 3.3 |
| A1_No_Mismatch | chr13 | 35,974,534 | 2 | 76 | 2.6 |
| A1_R2A | chr13 | 39,422,995 | 71 | 81 | 87.7 |
| A1_14G | chr13 | 45,660,680 | 8 | 91 | 8.8 |
| A1_18A | chr13 | 52,825,871 | 16 | 59 | 27.1 |
| A1_No_Mismatch | chr13 | 52,984,471 | 7 | 82 | 8.5 |
| A1_7C | chr13 | 54,390,323 | 5 | 50 | 10.0 |
| A1_No_Mismatch | chr13 | 58,195,963 | 2 | 49 | 4.1 |
| A1_No_Mismatch | chr13 | 59,198,980 | 4 | 74 | 5.4 |
| A1_No_Mismatch | chr13 | 60,553,748 | 3 | 59 | 5.1 |
| A1_7A | chr13 | 61,722,899 | 1 | 46 | 2.2 |
| A1_6G | chr13 | 63,393,220 | 5 | 32 | 15.6 |
| A1_No_Mismatch | chr13 | 63,557,971 | 4 | 65 | 6.2 |
| A1_No_Mismatch | chr13 | 81,167,064 | 3 | 49 | 6.1 |
| A1_R2A | chr13 | 84,989,960 | 36 | 40 | 90.0 |
| A1_No_Mismatch | chr13 | 89,849,514 | 3 | 48 | 6.3 |
| A1_No_Mismatch | chr13 | 90,827,212 | 2 | 87 | 2.3 |
| A1_No_Mismatch | chr13 | 91,239,399 | 2 | 21 | 9.5 |
| A1_No_Mismatch | chr13 | 91,427,845 | 3 | 69 | 4.3 |
| A1_No_Mismatch | chr13 | 92,359,799 | 6 | 54 | 11.1 |
| A1_16T | chr13 | 92,371,716 | 6 | 53 | 11.3 |
| A1_No_Mismatch | chr13 | 97,533,173 | 3 | 41 | 7.3 |
| A1_No_Mismatch | chr13 | 97,851,378 | 4 | 87 | 4.6 |
| A1_2T | chr13 | 98,335,431 | 2 | 58 | 3.4 |
| A1_No_Mismatch | chr13 | 102,450,876 | 3 | 72 | 4.2 |
| A1_6G | chr13 | 103,169,038 | 8 | 80 | 10.0 |

| | | | | | |
|---|---|---|---|---|---|
| A1_R2A | chr13 | 104,258,265 | 39 | 41 | 95.1 |
| A1_No_Mismatch | chr13 | 105,295,429 | 4 | 86 | 4.7 |
| A1_No_Mismatch | chr13 | 109,702,205 | 4 | 69 | 5.8 |
| A1_R2A | chr13 | 110,499,424 | 94 | 106 | 88.7 |
| A1_No_Mismatch | chr14 | 20,928,103 | 3 | 80 | 3.8 |
| A1_No_Mismatch | chr14 | 25,223,525 | 5 | 78 | 6.4 |
| A1_No_Mismatch | chr14 | 29,718,241 | 1 | 55 | 1.8 |
| A1_No_Mismatch | chr14 | 33,317,571 | 3 | 87 | 3.4 |
| A1_No_Mismatch | chr14 | 35,359,827 | 4 | 62 | 6.5 |
| A1_18C | chr14 | 37,474,426 | 8 | 47 | 17.0 |
| A1_16T | chr14 | 41,310,545 | 12 | 89 | 13.5 |
| A1_No_Mismatch | chr14 | 44,791,435 | 1 | 53 | 1.9 |
| A1_17T | chr14 | 47,214,060 | 11 | 48 | 22.9 |
| A1_R2A | chr14 | 48,867,754 | 50 | 59 | 84.7 |
| A1_No_Mismatch | chr14 | 49,561,634 | 5 | 76 | 6.6 |
| A1_No_Mismatch | chr14 | 51,185,322 | 4 | 103 | 3.9 |
| A1_4T | chr14 | 51,665,785 | 2 | 73 | 2.7 |
| A1_No_Mismatch | chr14 | 53,008,541 | 3 | 79 | 3.8 |
| A1_16G | chr14 | 53,312,787 | 6 | 25 | 24.0 |
| A1_7A | chr14 | 53,453,752 | 4 | 60 | 6.7 |
| A1_No_Mismatch | chr14 | 56,174,569 | 2 | 67 | 3.0 |
| A1_16G | chr14 | 63,681,373 | 4 | 58 | 6.9 |
| A1_No_Mismatch | chr14 | 65,087,934 | 0 | 60 | 0.0 |
| A1_2T | chr14 | 66,371,716 | 4 | 67 | 6.0 |
| A1_19T | chr14 | 67,969,084 | 9 | 52 | 17.3 |
| A1_4T | chr14 | 71,282,197 | 6 | 68 | 8.8 |
| A1_No_Mismatch | chr14 | 72,302,883 | 6 | 46 | 13.0 |
| A1_No_Mismatch | chr14 | 73,340,904 | 2 | 83 | 2.4 |
| A1_R2A | chr14 | 76,724,846 | 36 | 86 | 41.9 |
| A1_No_Mismatch | chr14 | 80,348,669 | 7 | 63 | 11.1 |
| A1_16T | chr14 | 90,090,747 | 8 | 71 | 11.3 |
| A1_R2A | chr14 | 98,440,426 | 77 | 87 | 88.5 |
| A1_4G | chr14 | 101,742,636 | 5 | 58 | 8.6 |
| A1_No_Mismatch | chr14 | 103,267,564 | 7 | 85 | 8.2 |
| A1_No_Mismatch | chr14 | 104,067,899 | 6 | 56 | 10.7 |
| A2_20C | chr1 | 3,010,899 | 59 | 77 | 76.6 |
| A2_20C | chr1 | 3,575,719 | 60 | 71 | 84.5 |
| A2_20T | chr1 | 6,976,875 | 26 | 76 | 34.2 |
| A2_No_mismatch | chr1 | 7,196,125 | 2 | 33 | 6.1 |
| A2_20C | chr1 | 9,020,839 | 85 | 97 | 87.6 |
| A2_20T | chr1 | 15,045,833 | 20 | 54 | 37.0 |

| | | | | | |
|---|---|---|---|---|---|
| A2_20T | chr1 | 17,188,836 | 60 | 82 | 73.2 |
| A2_20T | chr1 | 17,704,401 | 45 | 65 | 69.2 |
| A2_20G | chr1 | 20,132,516 | 21 | 44 | 47.7 |
| A2_20C | chr1 | 25,333,855 | 31 | 54 | 57.4 |
| A2_17A | chr1 | 27,805,053 | 1 | 50 | 2.0 |
| A2_No_mismatch | chr1 | 33,547,976 | 3 | 67 | 4.5 |
| A2_20T | chr1 | 40,505,890 | 10 | 64 | 15.6 |
| A2_20T | chr1 | 50,756,836 | 7 | 47 | 14.9 |
| A2_20T | chr1 | 53,341,255 | 20 | 62 | 32.3 |
| A2_No_mismatch | chr1 | 53,758,468 | 7 | 94 | 7.4 |
| A2_20T | chr1 | 57,938,449 | 71 | 98 | 72.4 |
| A2_20T | chr1 | 64,169,450 | 16 | 34 | 47.1 |
| A2_20G | chr1 | 73,213,067 | 26 | 37 | 70.3 |
| A2_R2A | chr1 | 87,503,036 | 44 | 60 | 73.3 |
| A2_20T | chr1 | 90,757,651 | 80 | 90 | 88.9 |
| A2_20T | chr1 | 90,862,992 | 42 | 55 | 76.4 |
| A2_20T | chr1 | 94,428,598 | 15 | 65 | 23.1 |
| A2_20C | chr1 | 99,116,782 | 34 | 49 | 69.4 |
| A2_20C | chr1 | 100,781,811 | 35 | 41 | 85.4 |
| A2_20T | chr1 | 101,979,427 | 50 | 75 | 66.7 |
| A2_20C | chr1 | 114,194,141 | 70 | 93 | 75.3 |
| A2_20T | chr1 | 157,674,504 | 31 | 51 | 60.8 |
| A2_20T | chr1 | 157,759,092 | 39 | 57 | 68.4 |
| A2_20T | chr1 | 161,754,685 | 48 | 82 | 58.5 |
| A2_20C | chr1 | 165,460,138 | 62 | 72 | 86.1 |
| A2_No_mismatch | chr1 | 167,597,385 | 5 | 74 | 6.8 |
| A2_No_mismatch | chr1 | 167,884,403 | 2 | 50 | 4.0 |
| A2_20C | chr1 | 170,834,489 | 33 | 42 | 78.6 |
| A2_20C | chr1 | 171,126,233 | 32 | 34 | 94.1 |
| A2_20T | chr1 | 179,302,404 | 59 | 60 | 98.3 |
| A2_20T | chr1 | 187,736,128 | 37 | 61 | 60.7 |
| A2_20T | chr1 | 189,905,471 | 58 | 66 | 87.9 |
| A2_20T | chr1 | 198,798,116 | 9 | 41 | 22.0 |
| A2_1T | chr1 | 199,678,208 | 5 | 67 | 7.5 |
| A2_20T | chr1 | 213,815,084 | 29 | 67 | 43.3 |
| A2_20T | chr1 | 213,824,621 | 45 | 66 | 68.2 |
| A2_20C | chr1 | 215,070,612 | 59 | 75 | 78.7 |
| A2_20T | chr1 | 222,380,320 | 23 | 67 | 34.3 |
| A2_17A | chr1 | 223,046,720 | 8 | 53 | 15.1 |
| A2_20T | chr1 | 230,360,527 | 29 | 69 | 42.0 |
| A2_20T | chr1 | 239,507,313 | 48 | 88 | 54.5 |

| | | | | | |
|---|---|---|---|---|---|
| A2_20T | chr1 | 240,751,949 | 61 | 75 | 81.3 |
| A2_20T | chr1 | 241,537,795 | 37 | 46 | 80.4 |
| A2_20T | chr1 | 244,309,326 | 62 | 81 | 76.5 |
| A2_20T | chr10 | 2,092,911 | 30 | 41 | 73.2 |
| A2_3T | chr10 | 3,358,047 | 10 | 52 | 19.2 |
| A2_3A | chr10 | 4,161,849 | 8 | 71 | 11.3 |
| A2_20T | chr10 | 17,526,197 | 35 | 47 | 74.5 |
| A2_20C | chr10 | 20,515,510 | 38 | 42 | 90.5 |
| A2_No_mismatch | chr10 | 21,246,975 | 2 | 96 | 2.1 |
| A2_20T | chr10 | 29,673,672 | 58 | 103 | 56.3 |
| A2_20T | chr10 | 30,756,810 | 28 | 87 | 32.2 |
| A2_No_mismatch | chr10 | 30,795,792 | 4 | 65 | 6.2 |
| A2_20T | chr10 | 32,222,853 | 21 | 52 | 40.4 |
| A2_20T | chr10 | 33,359,675 | 43 | 77 | 55.8 |
| A2_20T | chr10 | 52,674,365 | 62 | 73 | 84.9 |
| A2_20C | chr10 | 53,189,041 | 22 | 27 | 81.5 |
| A2_20T | chr10 | 59,053,761 | 24 | 40 | 60.0 |
| A2_20T | chr10 | 61,191,716 | 21 | 27 | 77.8 |
| A2_No_mismatch | chr10 | 66,116,908 | 3 | 84 | 3.6 |
| A2_20G | chr10 | 88,798,396 | 45 | 90 | 50.0 |
| A2_18A | chr10 | 109,596,331 | 3 | 52 | 5.8 |
| A2_4C | chr10 | 128,920,976 | 5 | 87 | 5.7 |
| A2_20C | chr11 | 26,241,050 | 57 | 69 | 82.6 |
| A2_6C | chr11 | 29,453,378 | 1 | 53 | 1.9 |
| A2_No_mismatch | chr11 | 77,853,494 | 3 | 61 | 4.9 |
| A2_18G | chr11 | 81,867,993 | 52 | 63 | 82.5 |
| A2_20C | chr11 | 107,395,160 | 45 | 64 | 70.3 |
| A2_6G | chr11 | 111,118,259 | 3 | 93 | 3.2 |
| A2_15T | chr11 | 121,371,008 | 8 | 68 | 11.8 |
| A2_No_mismatch | chr12 | 13,268,099 | 5 | 71 | 7.0 |
| A2_No_mismatch | chr12 | 28,173,252 | 3 | 33 | 9.1 |
| A2_16G | chr12 | 89,514,162 | 18 | 62 | 29.0 |
| A2_No_mismatch | chr12 | 102,675,999 | 4 | 30 | 13.3 |
| A2_No_mismatch | chr12 | 107,591,525 | 1 | 63 | 1.6 |
| A2_5G | chr12 | 113,027,845 | 4 | 84 | 4.8 |
| A2_No_mismatch | chr12 | 115,187,839 | 6 | 60 | 10.0 |
| A2_No_mismatch | chr13 | 35,974,533 | 1 | 73 | 1.4 |
| A2_13G | chr13 | 45,660,681 | 4 | 67 | 6.0 |
| A2_No_mismatch | chr13 | 52,984,472 | 5 | 84 | 6.0 |
| A2_14T | chr13 | 80,067,351 | 5 | 63 | 7.9 |
| A2_20C | chr13 | 103,993,845 | 39 | 47 | 83.0 |

| | | | | | |
|---|---|---|---|---|---|
| A2_No_mismatch | chr14 | 20,928,104 | 2 | 62 | 3.2 |
| A2_No_mismatch | chr14 | 25,223,526 | 4 | 80 | 5.0 |
| A2_No_mismatch | chr14 | 35,359,826 | 2 | 52 | 3.8 |
| A2_4C | chr14 | 46,956,845 | 3 | 49 | 6.1 |
| A2_No_mismatch | chr14 | 56,174,568 | 4 | 62 | 6.5 |
| A2_No_mismatch | chr14 | 103,267,565 | 4 | 96 | 4.2 |
| A2_No_mismatch | chr14 | 104,067,900 | 3 | 50 | 6.0 |
| A2_No_mismatch | chr15 | 26,606,071 | 8 | 79 | 10.1 |
| A2_5G | chr15 | 29,911,630 | 2 | 55 | 3.6 |
| A2_No_mismatch | chr15 | 63,172,455 | 2 | 85 | 2.4 |
| A2_No_mismatch | chr15 | 94,427,572 | 1 | 48 | 2.1 |
| A2_No_mismatch | chr16 | 31,347,584 | 2 | 60 | 3.3 |
| A2_No_mismatch | chr16 | 51,077,017 | 6 | 88 | 6.8 |
| A2_No_mismatch | chr16 | 71,487,230 | 3 | 84 | 3.6 |
| A2_No_mismatch | chr16 | 73,647,398 | 3 | 44 | 6.8 |
| A2_No_mismatch | chr16 | 73,647,400 | 2 | 44 | 4.5 |
| A2_5G | chr16 | 74,849,695 | 6 | 75 | 8.0 |
| A2_16G | chr17 | 63,505,532 | 42 | 67 | 62.7 |
| A2_15T | chr18 | 25,387,805 | 2 | 53 | 3.8 |
| A2_No_mismatch | chr18 | 33,259,624 | 0 | 36 | 0.0 |
| A2_5C | chr18 | 57,968,816 | 3 | 120 | 2.5 |
| A2_4C | chr2 | 6,855,234 | 3 | 87 | 3.4 |
| A2_No_mismatch | chr2 | 133,289,976 | 1 | 46 | 2.2 |
| A2_5G | chr20 | 18,629,898 | 0 | 38 | 0.0 |
| A2_No_mismatch | chr20 | 37,609,997 | 2 | 101 | 2.0 |
| A2_16G | chr20 | 47,203,426 | 31 | 98 | 31.6 |
| A2_1T | chr20 | 55,401,415 | 11 | 57 | 19.3 |
| A2_15G | chr21 | 20,524,885 | 8 | 34 | 23.5 |
| A2_6C | chr21 | 26,804,148 | 4 | 54 | 7.4 |
| A2_6G | chr21 | 30,427,249 | 0 | 98 | 0.0 |
| A2_15T | chr22 | 33,950,607 | 0 | 89 | 0.0 |
| A2_No_mismatch | chr23 | 5,304,359 | 2 | 15 | 13.3 |
| A2_15A | chr23 | 147,643,816 | 2 | 42 | 4.8 |
| A2_No_mismatch | chr3 | 101,248,224 | 2 | 34 | 5.9 |
| A2_16G | chr3 | 139,382,222 | 17 | 49 | 34.7 |
| A2_No_mismatch | chr3 | 159,301,614 | 7 | 66 | 10.6 |
| A2_3T | chr3 | 190,987,274 | 9 | 59 | 15.3 |
| A2_4G | chr4 | 33,233,155 | 1 | 45 | 2.2 |
| A2_17C | chr4 | 72,555,791 | 3 | 43 | 7.0 |
| A2_15G | chr4 | 83,384,247 | 3 | 26 | 11.5 |
| A2_No_mismatch | chr4 | 101,603,231 | 3 | 57 | 5.3 |

| | | | | | |
|---|---|---|---|---|---|
| A2_5G | chr4 | 132,365,768 | 4 | 66 | 6.1 |
| A2_13C | chr4 | 157,798,475 | 5 | 48 | 10.4 |
| A2_5T | chr4 | 185,834,995 | 2 | 53 | 3.8 |
| A2_6C | chr5 | 84,541,352 | 7 | 59 | 11.9 |
| A2_4C | chr5 | 131,614,547 | 4 | 67 | 6.0 |
| A2_15A | chr6 | 80,901,485 | 0 | 32 | 0.0 |
| A2_15G | chr6 | 104,611,533 | 7 | 55 | 12.7 |
| A2_No_mismatch | chr7 | 681,050 | 1 | 82 | 1.2 |
| A2_5C | chr7 | 4,574,391 | 6 | 95 | 6.3 |
| A2_1A | chr7 | 10,893,330 | 0 | 21 | 0.0 |
| A2_5G | chr7 | 19,354,895 | 1 | 35 | 2.9 |
| A2_13T | chr7 | 20,028,228 | 5 | 36 | 13.9 |
| A2_14G | chr8 | 15,341,534 | 7 | 87 | 8.0 |
| A2_4C | chr8 | 27,547,054 | 1 | 66 | 1.5 |
| A2_No_mismatch | chr8 | 34,238,820 | 4 | 47 | 8.5 |
| A2_5C | chr8 | 60,500,343 | 2 | 67 | 3.0 |
| A2_5G | chr9 | 83,483,149 | 2 | 95 | 2.1 |
| A2_R1A | chr9 | 114,416,464 | 59 | 81 | 72.8 |
| A5_No_Mismatch | chr1 | 3,010,897 | 1 | 99 | 1.0 |
| A5_No_Mismatch | chr1 | 3,575,717 | 4 | 65 | 6.2 |
| A5_20T | chr1 | 6,976,875 | 16 | 52 | 30.8 |
| A5_20A | chr1 | 7,196,127 | 32 | 47 | 68.1 |
| A5_No_Mismatch | chr1 | 9,020,860 | 3 | 100 | 3.0 |
| A5_20T | chr1 | 15,045,833 | 48 | 69 | 69.6 |
| A5_20T | chr1 | 17,188,836 | 87 | 103 | 84.5 |
| A5_20T | chr1 | 17,704,401 | 68 | 90 | 75.6 |
| A5_20G | chr1 | 20,132,516 | 28 | 64 | 43.8 |
| A5_No_Mismatch | chr1 | 25,333,853 | 3 | 91 | 3.3 |
| A5_20A | chr1 | 33,547,978 | 73 | 102 | 71.6 |
| A5_20T | chr1 | 40,505,890 | 45 | 73 | 61.6 |
| A5_15T | chr1 | 42,027,844 | 5 | 90 | 5.6 |
| A5_20A | chr1 | 47,596,160 | 63 | 83 | 75.9 |
| A5_20T | chr1 | 53,341,255 | 24 | 43 | 55.8 |
| A5_20A | chr1 | 53,758,447 | 75 | 106 | 70.8 |
| A5_17T | chr1 | 56,813,431 | 12 | 60 | 20.0 |
| A5_20T | chr1 | 57,938,449 | 90 | 113 | 79.6 |
| A5_20T | chr1 | 64,169,450 | 19 | 60 | 31.7 |
| A5_20G | chr1 | 73,213,067 | 42 | 46 | 91.3 |
| A5_15T | chr1 | 82,610,367 | 4 | 69 | 5.8 |
| A5_02T | chr1 | 84,568,733 | 13 | 66 | 19.7 |
| A5_20T | chr1 | 86,981,146 | 28 | 33 | 84.8 |

| | | | | | |
|---|---|---|---|---|---|
| A5_20T | chr1 | 90,757,651 | 101 | 117 | 86.3 |
| A5_20T | chr1 | 90,862,992 | 66 | 91 | 72.5 |
| A5_20T | chr1 | 94,428,598 | 21 | 82 | 25.6 |
| A5_No_Mismatch | chr1 | 99,116,780 | 9 | 93 | 9.7 |
| A5_No_Mismatch | chr1 | 100,781,809 | 2 | 48 | 4.2 |
| A5_20T | chr1 | 101,979,427 | 71 | 92 | 77.2 |
| A5_No_Mismatch | chr1 | 114,194,139 | 12 | 114 | 10.5 |
| A5_No_Mismatch | chr1 | 117,983,231 | 1 | 72 | 1.4 |
| A5_02C | chr1 | 118,375,666 | 6 | 44 | 13.6 |
| A5_20T | chr1 | 157,674,504 | 49 | 74 | 66.2 |
| A5_20T | chr1 | 157,759,092 | 61 | 82 | 74.4 |
| A5_15T | chr1 | 159,995,543 | 4 | 71 | 5.6 |
| A5_No_Mismatch | chr1 | 165,460,136 | 12 | 112 | 10.7 |
| A5_20A | chr1 | 167,597,364 | 26 | 78 | 33.3 |
| A5_20A | chr1 | 167,884,382 | 19 | 52 | 36.5 |
| A5_No_Mismatch | chr1 | 170,834,510 | 3 | 44 | 6.8 |
| A5_No_Mismatch | chr1 | 171,126,254 | 3 | 36 | 8.3 |
| A5_20T | chr1 | 187,736,128 | 42 | 92 | 45.7 |
| A5_20T | chr1 | 189,905,471 | 50 | 63 | 79.4 |
| A5_06C | chr1 | 191,076,859 | 2 | 58 | 3.4 |
| A5_R2A | chr1 | 198,355,521 | 75 | 98 | 76.5 |
| A5_20T | chr1 | 198,798,116 | 35 | 46 | 76.1 |
| A5_20T | chr1 | 213,815,084 | 61 | 84 | 72.6 |
| A5_20T | chr1 | 213,824,621 | 60 | 81 | 74.1 |
| A5_No_Mismatch | chr1 | 215,070,610 | 5 | 70 | 7.1 |
| A5_20T | chr1 | 222,380,320 | 33 | 67 | 49.3 |
| A5_05G | chr10 | 12,817,100 | 2 | 55 | 3.6 |
| A5_No_Mismatch | chr10 | 20,515,508 | 12 | 51 | 23.5 |
| A5_20A | chr10 | 21,246,977 | 85 | 121 | 70.2 |
| A5_20A | chr10 | 30,795,771 | 41 | 57 | 71.9 |
| A5_No_Mismatch | chr10 | 53,189,062 | 3 | 41 | 7.3 |
| A5_No_Mismatch | chr10 | 53,416,569 | 9 | 60 | 15.0 |
| A5_No_Mismatch | chr10 | 56,952,094 | 8 | 59 | 13.6 |
| A5_03G | chr10 | 60,132,369 | 1 | 59 | 1.7 |
| A5_06G | chr10 | 64,306,211 | 6 | 91 | 6.6 |
| A5_20A | chr10 | 66,116,887 | 55 | 79 | 69.6 |
| A5_20G | chr10 | 88,798,396 | 73 | 90 | 81.1 |
| A5_No_Mismatch | chr10 | 108,007,037 | 7 | 103 | 6.8 |
| A5_15T | chr10 | 115,106,616 | 7 | 93 | 7.5 |
| A5_15T | chr10 | 117,302,753 | 9 | 115 | 7.8 |
| A5_No_Mismatch | chr10 | 117,453,762 | 6 | 103 | 5.8 |

| | | | | | |
|---|---|---|---|---|---|
| A5_04C | chr10 | 117,809,629 | 5 | 55 | 9.1 |
| A5_17A | chr10 | 123,018,828 | 5 | 103 | 4.9 |
| A5_20A | chr10 | 130,075,712 | 46 | 54 | 85.2 |
| A5_18A | chr11 | 13,657,741 | 10 | 91 | 11.0 |
| A5_No_Mismatch | chr11 | 14,681,409 | 2 | 61 | 3.3 |
| A5_R2A | chr11 | 23,663,455 | 84 | 87 | 96.6 |
| A5_No_Mismatch | chr11 | 26,241,071 | 9 | 65 | 13.8 |
| A5_15T | chr11 | 36,338,930 | 8 | 78 | 10.3 |
| A5_05G | chr11 | 45,413,654 | 3 | 60 | 5.0 |
| A5_15T | chr11 | 57,168,861 | 6 | 97 | 6.2 |
| A5_No_Mismatch | chr11 | 70,899,262 | 8 | 81 | 9.9 |
| A5_14G | chr11 | 77,843,969 | 5 | 74 | 6.8 |
| A5_No_Mismatch | chr11 | 92,723,798 | 3 | 61 | 4.9 |
| A5_No_Mismatch | chr11 | 101,318,613 | 3 | 39 | 7.7 |
| A5_15T | chr11 | 102,126,423 | 5 | 81 | 6.2 |
| A5_No_Mismatch | chr11 | 103,711,097 | 5 | 98 | 5.1 |
| A5_No_Mismatch | chr11 | 107,395,181 | 6 | 61 | 9.8 |
| A5_No_Mismatch | chr11 | 115,358,478 | 6 | 67 | 9.0 |
| A5_No_Mismatch | chr11 | 122,805,626 | 4 | 78 | 5.1 |
| A5_14G | chr11 | 123,774,594 | 2 | 98 | 2.0 |
| A5_15T | chr11 | 125,839,750 | 6 | 64 | 9.4 |
| A5_NG | chr12 | 10,536,888 | 3 | 64 | 4.7 |
| A5_NA | chr12 | 20,286,094 | 7 | 60 | 11.7 |
| A5_17A | chr12 | 21,883,331 | 5 | 66 | 7.6 |
| A5_15T | chr12 | 23,746,874 | 2 | 51 | 3.9 |
| A5_15T | chr12 | 87,182,060 | 12 | 67 | 17.9 |
| A5_NA | chr12 | 92,061,992 | 2 | 15 | 13.3 |
| A5_R1A | chr12 | 95,920,210 | 55 | 70 | 78.6 |
| A5_NT | chr12 | 98,920,575 | 5 | 92 | 5.4 |
| A5_15T | chr12 | 102,742,775 | 6 | 64 | 9.4 |
| A5_05G | chr12 | 125,322,185 | 6 | 68 | 8.8 |
| A5_18T | chr13 | 42,211,877 | 24 | 62 | 38.7 |
| A5_15T | chr13 | 53,164,687 | 9 | 97 | 9.3 |
| A5_15T | chr13 | 57,847,068 | 16 | 53 | 30.2 |
| A5_05G | chr13 | 60,512,247 | 2 | 45 | 4.4 |
| A5_05C | chr13 | 61,683,448 | 7 | 57 | 12.3 |
| A5_06A | chr13 | 61,722,898 | 3 | 64 | 4.7 |
| A5_05G | chr13 | 62,914,941 | 6 | 48 | 12.5 |
| A5_15T | chr13 | 87,332,279 | 6 | 70.0 | 8.6 |
| A5_No_Mismatch | chr13 | 103,993,866 | 4 | 56 | 7.1 |
| A5_03G | chr14 | 29,124,615 | 10 | 65 | 15.4 |

| | | | | | |
|---|---|---|---|---|---|
| A5_15T | chr14 | 41,098,568 | 2 | 35 | 5.7 |
| A5_No_Mismatch | chr14 | 53,754,073 | 1 | 24 | 4.2 |
| A5_No_Mismatch | chr14 | 64,872,804 | 3 | 69 | 4.3 |
| A5_04C | chr14 | 82,934,864 | 0 | 78 | 0.0 |
| A5_No_Mismatch | chr14 | 88,966,311 | 6 | 121 | 5.0 |
| A5_No_Mismatch | chr14 | 101,413,512 | 3 | 72 | 4.2 |
| A5_No_Mismatch | chr15 | 37,367,340 | 1 | 77 | 1.3 |
| A5_15T | chr15 | 43,197,889 | 7 | 56 | 12.5 |
| A5_No_Mismatch | chr15 | 73,182,145 | 1 | 57 | 1.8 |
| A5_No_Mismatch | chr15 | 84,966,752 | 7 | 76 | 9.2 |
| A5_No_Mismatch | chr15 | 88,971,443 | 2 | 133 | 1.5 |
| A5_No_Mismatch | chr15 | 91,896,552 | 2 | 95 | 2.1 |
| A5_15T | chr15 | 101,417,106 | 12 | 114 | 10.5 |
| A5_15A | chr16 | 10,671,851 | 10 | 102 | 9.8 |
| A5_No_Mismatch | chr16 | 15,276,473 | 5 | 92 | 5.4 |
| A5_No_Mismatch | chr16 | 17,898,682 | 3 | 93 | 3.2 |
| A5_20A | chr16 | 73,647,400 | 60 | 81 | 74.1 |
| A5_04A | chr16 | 75,885,051 | 9 | 110 | 8.2 |
| A5_15T | chr16 | 76,309,774 | 5 | 59 | 8.5 |
| A5_No_Mismatch | chr16 | 76,958,205 | 5 | 55 | 9.1 |
| A5_04C | chr17 | 56,731,246 | 3 | 128 | 2.3 |
| A5_No_Mismatch | chr17 | 72,036,657 | 3 | 91 | 3.3 |
| A5_15T | chr17 | 72,692,575 | 2 | 57 | 3.5 |
| A5_15G | chr17 | 76,113,323 | 4 | 77 | 5.2 |
| A5_No_Mismatch | chr17 | 76,852,991 | 7 | 91 | 7.7 |
| A5_14G | chr17 | 79,062,877 | 19 | 86 | 22.1 |
| A5_No_Mismatch | chr18 | 1,418,517 | 10 | 93 | 10.8 |
| A5_15T | chr18 | 31,140,927 | 4 | 53 | 7.5 |
| A5_15T | chr18 | 47,976,206 | 5 | 32 | 15.6 |
| A5_No_Mismatch | chr18 | 63,572,691 | 8 | 100 | 8.0 |
| A5_15T | chr18 | 63,761,659 | 5 | 99 | 5.1 |
| A5_No_Mismatch | chr18 | 72,712,613 | 3 | 81 | 3.7 |
| A5_15T | chr18 | 74,567,057 | 10 | 81 | 12.3 |
| A5_No_Mismatch | chr19 | 29,423,901 | 4 | 62 | 6.5 |
| A5_No_Mismatch | chr19 | 33,714,730 | 5 | 99 | 5.1 |
| A5_No_Mismatch | chr2 | 3,721,772 | 2 | 56 | 3.6 |
| A5_R1A | chr2 | 17,072,049 | 45 | 61 | 73.8 |
| A5_No_Mismatch | chr2 | 32,983,662 | 5 | 73 | 6.8 |
| A5_16T | chr2 | 37,900,000 | 11 | 98 | 11.2 |
| A5_No_Mismatch | chr2 | 41,862,146 | 6 | 116 | 5.2 |
| A5_No_Mismatch | chr2 | 58,804,349 | 17 | 54 | 31.5 |

| | | | | | |
|---|---|---|---|---|---|
| A5_R1C | chr2 | 81,794,515 | 104 | 110 | 94.5 |
| A5_No_Mismatch | chr2 | 98,667,039 | 3 | 88 | 3.4 |
| A5_05G | chr2 | 122,411,673 | 9 | 72 | 12.5 |
| A5_R1A | chr2 | 129,020,245 | 62 | 88 | 70.5 |
| A5_No_Mismatch | chr2 | 143,815,826 | 23 | 65 | 35.4 |
| A5_No_Mismatch | chr2 | 153,589,762 | 3 | 30 | 10.0 |
| A5_No_Mismatch | chr2 | 192,913,994 | 3 | 54 | 5.6 |
| A5_No_Mismatch | chr2 | 205,419,218 | 3 | 52 | 5.8 |
| A5_02T | chr2 | 224,397,194 | 3 | 58 | 5.2 |
| A5_01T | chr2 | 225,205,629 | 9 | 71 | 12.7 |
| A5_No_Mismatch | chr2 | 235,154,641 | 5 | 72 | 6.9 |
| A5_05T | chr20 | 37,664,877 | 2 | 98 | 2.0 |
| A5_No_Mismatch | chr20 | 48,775,685 | 0 | 23 | 0.0 |
| A5_No_Mismatch | chr20 | 51,062,160 | 2 | 119 | 1.7 |
| A5_05G | chr20 | 52,482,548 | 7 | 85 | 8.2 |
| A5_06C | chr21 | 30,698,871 | 0 | 19 | 0.0 |
| A5_19G | chr21 | 35,296,864 | 78 | 84 | 92.9 |
| A5_No_Mismatch | chr21 | 44,776,666 | 2 | 43 | 4.7 |
| A5_No_Mismatch | chr22 | 34,008,067 | 2 | 72 | 2.8 |
| A5_05C | chr22 | 47,493,086 | 6 | 93 | 6.5 |
| A5_15G | chr23 | 5,539,274 | 1 | 31 | 3.2 |
| A5_No_Mismatch | chr23 | 7,125,822 | 2 | 58 | 3.4 |
| A5_R1A | chr23 | 23,632,280 | 7 | 27 | 25.9 |
| A5_No_Mismatch | chr23 | 55,385,471 | 1 | 44 | 2.3 |
| A5_R2A | chr23 | 90,687,686 | 33 | 38 | 86.8 |
| A5_05C | chr23 | 139,771,169 | 1 | 18 | 5.6 |
| A5_20A | chr3 | 159,301,614 | 61 | 70 | 87.1 |
| A5_01T | chr3 | 171,053,725 | 7 | 81 | 8.6 |
| A5_20A | chr3 | 171,488,780 | 56 | 73 | 76.7 |
| A5_15T | chr3 | 176,459,125 | 5 | 63 | 7.9 |
| A5_No_Mismatch | chr3 | 183,484,601 | 36 | 74 | 48.6 |
| A5_19G | chr4 | 39,331,970 | 50 | 55 | 90.9 |
| A5_17T | chr4 | 42,463,307 | 5 | 78 | 6.4 |
| A5_05G | chr4 | 52,548,820 | 2 | 104 | 1.9 |
| A5_04C | chr4 | 78,838,214 | 5 | 53 | 9.4 |
| A5_13G | chr4 | 155,387,555 | 6 | 108 | 5.6 |
| A5_06A | chr4 | 157,723,000 | 7 | 53 | 13.2 |
| A5_15T | chr5 | 40,881,610 | 7 | 97 | 7.2 |
| A5_No_Mismatch | chr5 | 110,975,458 | 6 | 70 | 8.6 |
| A5_R2A | chr5 | 151,958,789 | 80 | 95 | 84.2 |
| A5_19G | chr5 | 155,430,584 | 98 | 104 | 94.2 |

| | | | | | |
|---|---|---|---|---|---|
| A5_No_Mismatch | chr6 | 67,578,212 | 35 | 48 | 72.9 |
| A5_13G | chr6 | 68,261,835 | 40 | 54 | 74.1 |
| A5_18T | chr6 | 77,202,024 | 1 | 17 | 5.9 |
| A5_No_Mismatch | chr6 | 107,306,305 | 3 | 84 | 3.6 |
| A5_R1A | chr6 | 134,150,581 | 42 | 51 | 82.4 |
| A5_15T | chr6 | 134,152,515 | 3 | 55 | 5.5 |
| A5_No_Mismatch | chr6 | 137,833,937 | 6 | 75 | 8.0 |
| A5_05C | chr7 | 4,243,637 | 1 | 72 | 1.4 |
| A5_15T | chr7 | 29,945,341 | 8 | 100 | 8.0 |
| A5_14G | chr7 | 70,350,554 | 7 | 63 | 11.1 |
| A5_05C | chr7 | 89,076,311 | 1 | 60 | 1.7 |
| A5_15T | chr7 | 116,046,361 | 4 | 41 | 9.8 |
| A5_05G | chr7 | 123,032,810 | 4 | 62 | 6.5 |
| A5_No_Mismatch | chr7 | 133,223,754 | 4 | 60 | 6.7 |
| A5_No_Mismatch | chr8 | 109,011,651 | 3 | 49 | 6.1 |
| A5_R2C | chr8 | 110,581,946 | 93 | 96 | 96.9 |
| A5_17A | chr9 | 14,142,246 | 3 | 74 | 4.1 |
| A5_01G | chr9 | 110,227,359 | 3 | 76 | 3.9 |
| A5_15T | chr9 | 110,320,301 | 0 | 78 | 0.0 |
| A5_15T | chr9 | 119,157,140 | 4 | 39 | 10.3 |
| A7_20C | chr1 | 3,010,899 | 93 | 101 | 92.1 |
| A7_20C | chr1 | 3,575,719 | 51 | 59 | 86.4 |
| A7_05G | chr1 | 4,099,320 | 2 | 76 | 2.6 |
| A7_R1A | chr1 | 4,669,531 | 48 | 78 | 61.5 |
| A7_No_Mismatch | chr1 | 6,976,873 | 1 | 74 | 1.4 |
| A7_20A | chr1 | 7,196,127 | 29 | 34 | 85.3 |
| A7_20C | chr1 | 9,020,839 | 40 | 62 | 64.5 |
| A7_R2A | chr1 | 11,113,897 | 33 | 52 | 63.5 |
| A7_03T | chr1 | 13,847,586 | 1 | 74 | 1.4 |
| A7_18T | chr1 | 14,428,767 | 3 | 55 | 5.5 |
| A7_No_Mismatch | chr1 | 15,045,854 | 5 | 75 | 6.7 |
| A7_No_Mismatch | chr1 | 17,704,399 | 4 | 85 | 4.7 |
| A7_20G | chr1 | 20,132,516 | 26 | 48 | 54.2 |
| A7_20C | chr1 | 25,333,855 | 17 | 70 | 24.3 |
| A7_01T | chr1 | 33,239,678 | 6 | 108 | 5.6 |
| A7_20A | chr1 | 33,547,978 | 73 | 99 | 73.7 |
| A7_No_Mismatch | chr1 | 40,505,888 | 1 | 95 | 1.1 |
| A7_No_Mismatch | chr1 | 53,341,253 | 0 | 78 | 0.0 |
| A7_20A | chr1 | 53,758,447 | 76 | 110 | 69.1 |
| A7_R2A | chr1 | 55,621,250 | 83 | 98 | 84.7 |
| A7_No_Mismatch | chr1 | 57,938,447 | 6 | 110 | 5.5 |

| | | | | | |
|---|---|---|---|---|---|
| A7_03G | chr1 | 58,513,154 | 0 | 36 | 0.0 |
| A7_R2A | chr1 | 59,031,477 | 74 | 91 | 81.3 |
| A7_06C | chr1 | 59,378,272 | 3 | 70 | 4.3 |
| A7_R2A | chr1 | 63,977,117 | 32 | 36 | 88.9 |
| A7_15T | chr1 | 67,509,308 | 9 | 55 | 16.4 |
| A7_R2C | chr1 | 69,605,159 | 48 | 52 | 92.3 |
| A7_15A | chr1 | 71,854,788 | 5 | 47 | 10.6 |
| A7_R2A | chr1 | 76,097,360 | 72 | 86 | 83.7 |
| A7_19C | chr1 | 78,122,602 | 60 | 74 | 81.1 |
| A7_15A | chr1 | 79,251,487 | 3 | 47 | 6.4 |
| A7_05G | chr1 | 83,141,838 | 8 | 104 | 7.7 |
| A7_20A | chr1 | 83,418,249 | 56 | 61 | 91.8 |
| A7_R1A | chr1 | 85,804,679 | 12 | 61 | 19.7 |
| A7_R1A | chr1 | 90,607,426 | 45 | 61 | 73.8 |
| A7_No_Mismatch | chr1 | 90,757,649 | 6 | 87 | 6.9 |
| A7_No_Mismatch | chr1 | 90,863,013 | 2 | 68 | 2.9 |
| A7_No_Mismatch | chr1 | 94,428,596 | 8 | 76 | 10.5 |
| A7_05G | chr1 | 96,760,017 | 5 | 129 | 3.9 |
| A7_18T | chr1 | 96,881,350 | 7 | 31 | 22.6 |
| A7_20C | chr1 | 99,116,782 | 58 | 68 | 85.3 |
| A7_20C | chr1 | 100,781,811 | 57 | 68 | 83.8 |
| A7_No_Mismatch | chr1 | 101,979,425 | 14 | 94 | 14.9 |
| A7_20C | chr1 | 114,194,141 | 84 | 105 | 80.0 |
| A7_04A | chr1 | 146,694,256 | 16 | 95 | 16.8 |
| A7_06C | chr1 | 151,868,314 | 10 | 94 | 10.6 |
| A7_No_Mismatch | chr1 | 157,674,502 | 7 | 81 | 8.6 |
| A7_No_Mismatch | chr1 | 157,759,090 | 5 | 91 | 5.5 |
| A7_No_Mismatch | chr1 | 161,754,706 | 12 | 111 | 10.8 |
| A7_20C | chr1 | 165,460,138 | 93 | 102 | 91.2 |
| A7_20A | chr1 | 167,597,364 | 46 | 67 | 68.7 |
| A7_20A | chr1 | 167,884,382 | 15 | 50 | 30.0 |
| A7_R2A | chr1 | 168,464,431 | 28 | 34 | 82.4 |
| A7_R1A | chr1 | 169,757,665 | 26 | 63 | 41.3 |
| A7_20C | chr1 | 170,834,489 | 39 | 40 | 97.5 |
| A7_20C | chr1 | 171,126,233 | 39 | 42 | 92.9 |
| A7_R1A | chr1 | 171,955,957 | 12 | 70 | 17.1 |
| A7_04C | chr1 | 179,146,658 | 5 | 107 | 4.7 |
| A7_R1A | chr1 | 187,507,308 | 9 | 32 | 28.1 |
| A7_No_Mismatch | chr1 | 187,736,149 | 6 | 67 | 9.0 |
| A7_No_Mismatch | chr1 | 189,905,492 | 6 | 56 | 10.7 |
| A7_15T | chr1 | 194,265,065 | 4 | 61 | 6.6 |

| | | | | | |
|---|---|---|---|---|---|
| A7_15T | chr1 | 196,670,470 | 5 | 58 | 8.6 |
| A7_No_Mismatch | chr1 | 198,798,137 | 3 | 50 | 6.0 |
| A7_15T | chr1 | 207,118,551 | 4 | 78 | 5.1 |
| A7_20C | chr1 | 215,070,612 | 70 | 79 | 88.6 |
| A7_01A | chr1 | 218,448,628 | 5 | 55 | 9.1 |
| A7_No_Mismatch | chr1 | 222,380,341 | 2 | 62 | 3.2 |
| A7_No_Mismatch | chr1 | 230,360,525 | 3 | 92 | 3.3 |
| A7_R1A | chr1 | 233,889,312 | 7 | 49 | 14.3 |
| A7_06C | chr1 | 237,616,937 | 4 | 84 | 4.8 |
| A7_No_Mismatch | chr1 | 239,507,311 | 10 | 107 | 9.3 |
| A7_No_Mismatch | chr1 | 240,751,947 | 9 | 95 | 9.5 |
| A7_No_Mismatch | chr1 | 241,537,816 | 23 | 48 | 47.9 |
| A7_No_Mismatch | chr1 | 244,309,324 | 4 | 90 | 4.4 |
| A7_R1A | chr10 | 2,902,169 | 9 | 58 | 15.5 |
| A7_14G | chr10 | 7,418,679 | 6 | 67 | 9.0 |
| A7_No_Mismatch | chr10 | 17,526,195 | 5 | 70 | 7.1 |
| A7_15T | chr10 | 17,786,881 | 7 | 90 | 7.8 |
| A7_20C | chr10 | 20,515,510 | 56 | 60 | 93.3 |
| A7_20A | chr10 | 21,246,977 | 62 | 139 | 44.6 |
| A7_05G | chr10 | 22,032,393 | 5 | 56 | 8.9 |
| A7_15T | chr10 | 23,489,229 | 6 | 52 | 11.5 |
| A7_No_Mismatch | chr10 | 29,673,693 | 4 | 127 | 3.1 |
| A7_No_Mismatch | chr10 | 30,756,831 | 3 | 79 | 3.8 |
| A7_20A | chr10 | 30,795,771 | 48 | 60 | 80.0 |
| A7_No_Mismatch | chr10 | 32,222,874 | 2 | 60 | 3.3 |
| A7_No_Mismatch | chr10 | 33,359,696 | 5 | 72 | 6.9 |
| A7_15A | chr10 | 33,599,550 | 5 | 70 | 7.1 |
| A7_R1C | chr10 | 36,856,779 | 40 | 53 | 75.5 |
| A7_No_Mismatch | chr10 | 52,674,363 | 3 | 60 | 5.0 |
| A7_20C | chr10 | 53,189,041 | 19 | 19 | 100.0 |
| A7_R2A | chr10 | 56,065,058 | 55 | 62 | 88.7 |
| A7_18T | chr10 | 56,210,540 | 3 | 30 | 10.0 |
| A7_20C | chr10 | 56,952,096 | 54 | 64 | 84.4 |
| A7_18G | chr10 | 58,173,557 | 44 | 53 | 83.0 |
| A7_No_Mismatch | chr10 | 59,053,782 | 2 | 44 | 4.5 |
| A7_14G | chr10 | 59,278,145 | 6 | 74 | 8.1 |
| A7_No_Mismatch | chr10 | 61,126,798 | 2 | 75 | 2.7 |
| A7_20A | chr10 | 66,116,887 | 72 | 98 | 73.5 |
| A7_05G | chr10 | 75,376,864 | 3 | 78 | 3.8 |
| A7_15G | chr10 | 76,110,619 | 13 | 82 | 15.9 |
| A7_15T | chr10 | 78,242,162 | 16 | 128 | 12.5 |

| | | | | | |
|---|---|---|---|---|---|
| A7_No_Mismatch | chr10 | 82,339,505 | 5 | 70 | 7.1 |
| A7_20G | chr10 | 88,798,396 | 60 | 79 | 75.9 |
| A7_No_Mismatch | chr10 | 97,167,771 | 3 | 74 | 4.1 |
| A7_No_Mismatch | chr10 | 107,825,296 | 7 | 93 | 7.5 |
| A7_17A | chr10 | 108,185,355 | 11 | 66 | 16.7 |
| A7_18T | chr10 | 114,094,081 | 2 | 84 | 2.4 |
| A7_15T | chr10 | 118,855,899 | 8 | 92 | 8.7 |
| A7_No_Mismatch | chr10 | 124,369,573 | 6 | 59 | 10.2 |
| A7_18T | chr10 | 127,679,144 | 5 | 64 | 7.8 |
| A7_04C | chr10 | 129,072,161 | 6 | 90 | 6.7 |
| A7_15T | chr10 | 132,600,851 | 11 | 86 | 12.8 |
| A7_No_Mismatch | chr11 | 2,324,555 | 17 | 70 | 24.3 |
| A7_No_Mismatch | chr11 | 9,190,416 | 3 | 82 | 3.7 |
| A7_02T | chr11 | 12,022,286 | 3 | 54 | 5.6 |
| A7_No_Mismatch | chr11 | 15,499,747 | 6 | 62 | 9.7 |
| A7_15G | chr11 | 22,930,963 | 3 | 62 | 4.8 |
| A7_No_Mismatch | chr11 | 23,384,178 | 5 | 47 | 10.6 |
| A7_No_Mismatch | chr11 | 26,205,129 | 4 | 48 | 8.3 |
| A7_20C | chr11 | 26,241,050 | 59 | 62 | 95.2 |
| A7_No_Mismatch | chr11 | 29,441,740 | 4 | 54 | 7.4 |
| A7_No_Mismatch | chr11 | 38,748,214 | 7 | 68 | 10.3 |
| A7_No_Mismatch | chr11 | 40,625,503 | 1 | 13 | 7.7 |
| A7_15T | chr11 | 43,477,609 | 10 | 78 | 12.8 |
| A7_No_Mismatch | chr11 | 46,187,046 | 0 | 45 | 0.0 |
| A7_No_Mismatch | chr11 | 48,506,988 | 3 | 55 | 5.5 |
| A7_15T | chr11 | 50,141,888 | 4 | 77 | 5.2 |
| A7_15T | chr11 | 50,336,628 | 5 | 60 | 8.3 |
| A7_No_Mismatch | chr11 | 54,677,537 | 2 | 18 | 11.1 |
| A7_No_Mismatch | chr11 | 56,955,109 | 3 | 70 | 4.3 |
| A7_05G | chr11 | 61,153,218 | 4 | 78 | 5.1 |
| A7_No_Mismatch | chr11 | 74,826,390 | 7 | 38 | 18.4 |
| A7_15A | chr11 | 81,840,068 | 6 | 91 | 6.6 |
| A7_No_Mismatch | chr11 | 87,829,589 | 8 | 71 | 11.3 |
| A7_No_Mismatch | chr11 | 91,367,597 | 5 | 76 | 6.6 |
| A7_20C | chr11 | 107,395,160 | 52 | 69 | 75.4 |
| A7_13G | chr11 | 111,402,110 | 2 | 69 | 2.9 |
| A7_No_Mismatch | chr11 | 115,026,554 | 4 | 65 | 6.2 |
| A7_No_Mismatch | chr11 | 116,390,323 | 8 | 101 | 7.9 |
| A7_18T | chr11 | 117,397,732 | 5 | 80 | 6.3 |
| A7_18T | chr11 | 119,559,017 | 4 | 64 | 6.3 |
| A7_No_Mismatch | chr11 | 127,322,009 | 6 | 90 | 6.7 |

| | | | | | |
|---|---|---|---|---|---|
| A7_15T | chr11 | 132,508,623 | 3 | 84 | 3.6 |
| A7_15T | chr12 | 3,774,826 | 25 | 56 | 44.6 |
| A7_06C | chr12 | 8,166,433 | 3 | 49 | 6.1 |
| A7_17A | chr12 | 8,345,538 | 8 | 86 | 9.3 |
| A7_03A | chr12 | 31,951,338 | 56 | 73 | 76.7 |
| A7_No_Mismatch | chr12 | 38,673,072 | 4 | 52 | 7.7 |
| A7_18T | chr12 | 40,581,080 | 10 | 61 | 16.4 |
| A7_05G | chr12 | 47,401,346 | 3 | 71 | 4.2 |
| A7_No_Mismatch | chr12 | 58,355,079 | 1 | 63 | 1.6 |
| A7_No_Mismatch | chr12 | 58,859,975 | 3 | 82 | 3.7 |
| A7_17A | chr12 | 61,899,569 | 7 | 107 | 6.5 |
| A7_15T | chr12 | 65,920,587 | 4 | 62 | 6.5 |
| A7_15T | chr12 | 66,511,117 | 1 | 42 | 2.4 |
| A7_17A | chr12 | 72,229,367 | 4 | 77 | 5.2 |
| A7_03T | chr12 | 74,025,066 | 1 | 48 | 2.1 |
| A7_16T | chr12 | 82,620,919 | 0 | 16 | 0.0 |
| A7_04C | chr12 | 82,999,698 | 11 | 85 | 12.9 |
| A7_No_Mismatch | chr12 | 88,019,203 | 2 | 55 | 3.6 |
| A7_No_Mismatch | chr12 | 93,045,786 | 1 | 56 | 1.8 |
| A7_No_Mismatch | chr12 | 95,808,269 | 11 | 75 | 14.7 |
| A7_No_Mismatch | chr12 | 101,856,151 | 8 | 74 | 10.8 |
| A7_No_Mismatch | chr12 | 103,555,396 | 2 | 88 | 2.3 |
| A7_15T | chr12 | 104,396,082 | 4 | 52 | 7.7 |
| A7_No_Mismatch | chr12 | 105,927,627 | 6 | 57 | 10.5 |
| A7_15T | chr12 | 109,639,247 | 9 | 73 | 12.3 |
| A7_R1A | chr12 | 116,514,045 | 66 | 96 | 68.8 |
| A7_03T | chr12 | 122,160,862 | 2 | 79 | 2.5 |
| A7_03G | chr12 | 123,844,205 | 4 | 57 | 7.0 |
| A7_18T | chr12 | 130,063,533 | 10 | 104 | 9.6 |
| A7_16G | chr12 | 131,436,432 | 19 | 68 | 27.9 |
| A7_No_Mismatch | chr13 | 35,963,705 | 4 | 96 | 4.2 |
| A7_03T | chr13 | 45,205,322 | 4 | 59 | 6.8 |
| A7_17A | chr13 | 52,825,870 | 16 | 73 | 21.9 |
| A7_06C | chr13 | 54,390,322 | 2 | 62 | 3.2 |
| A7_17C | chr13 | 54,767,629 | 2 | 53 | 3.8 |
| A7_No_Mismatch | chr13 | 59,198,981 | 2 | 96 | 2.1 |
| A7_No_Mismatch | chr13 | 60,553,749 | 1 | 51 | 2.0 |
| A7_05G | chr13 | 63,393,219 | 6 | 41 | 14.6 |
| A7_No_Mismatch | chr13 | 63,557,970 | 2 | 72 | 2.8 |
| A7_No_Mismatch | chr13 | 81,167,065 | 0 | 72 | 0.0 |
| A7_No_Mismatch | chr13 | 82,052,880 | 6 | 54 | 11.1 |

| | | | | | |
|---|---|---|---|---|---|
| A7_18T | chr13 | 85,775,374 | 12 | 104 | 11.5 |
| A7_No_Mismatch | chr13 | 87,917,381 | 6 | 19 | 31.6 |
| A7_No_Mismatch | chr13 | 90,827,213 | 6 | 107 | 5.6 |
| A7_No_Mismatch | chr13 | 97,533,174 | 20 | 57 | 35.1 |
| A7_No_Mismatch | chr13 | 97,851,379 | 1 | 113 | 0.9 |
| A7_No_Mismatch | chr13 | 102,450,875 | 3 | 80 | 3.8 |
| A7_No_Mismatch | chr13 | 109,702,206 | 7 | 92 | 7.6 |
| A7_15T | chr14 | 26,245,653 | 3 | 13 | 23.1 |
| A7_No_Mismatch | chr14 | 29,718,240 | 4 | 56 | 7.1 |
| A7_15T | chr14 | 41,310,544 | 6 | 64 | 9.4 |
| A7_No_Mismatch | chr14 | 44,791,436 | 3 | 57 | 5.3 |
| A7_No_Mismatch | chr14 | 51,185,323 | 4 | 98 | 4.1 |
| A7_No_Mismatch | chr14 | 53,008,540 | 0 | 96 | 0.0 |
| A7_15G | chr14 | 53,312,788 | 4 | 29 | 13.8 |
| A7_05G | chr14 | 55,796,744 | 3 | 104 | 2.9 |
| A7_15G | chr14 | 63,681,372 | 6 | 58 | 10.3 |
| A7_No_Mismatch | chr14 | 65,087,933 | 8 | 69 | 11.6 |
| A7_No_Mismatch | chr14 | 72,302,882 | 6 | 57 | 10.5 |
| A7_No_Mismatch | chr14 | 73,340,903 | 9 | 81 | 11.1 |
| A7_15T | chr14 | 87,464,076 | 5 | 67 | 7.5 |
| A7_14T | chr14 | 88,118,882 | 9 | 57 | 15.8 |
| A7_15T | chr14 | 90,090,746 | 5 | 68 | 7.4 |
| A7_03G | chr15 | 23,765,770 | 0 | 73 | 0.0 |
| A7_05G | chr15 | 25,762,068 | 4 | 95 | 4.2 |
| A7_06G | chr15 | 48,694,587 | 6 | 61 | 9.8 |
| A7_No_Mismatch | chr15 | 52,724,572 | 5 | 62 | 8.1 |
| A7_05G | chr15 | 76,862,002 | 0 | 38 | 0.0 |
| A7_04A | chr15 | 91,698,020 | 5 | 89 | 5.6 |
| A7_05G | chr15 | 94,303,606 | 8 | 61 | 13.1 |
| A7_17A | chr15 | 95,994,158 | 3 | 107 | 2.8 |
| A7_04C | chr15 | 99,184,929 | 7 | 66 | 10.6 |
| A7_No_Mismatch | chr16 | 15,458,016 | 1 | 44 | 2.3 |
| A7_05G | chr16 | 25,563,836 | 5 | 58 | 8.6 |
| A7_06G | chr16 | 26,262,488 | 4 | 53 | 7.5 |
| A7_05G | chr16 | 26,927,147 | 12 | 83 | 14.5 |
| A7_17A | chr16 | 54,155,857 | 6 | 54 | 11.1 |
| A7_15T | chr16 | 57,725,273 | 4 | 90 | 4.4 |
| A7_05C | chr16 | 58,701,172 | 7 | 56 | 12.5 |
| A7_05G | chr16 | 58,937,938 | 5 | 78 | 6.4 |
| A7_17T | chr16 | 64,058,606 | 14 | 74 | 18.9 |
| A7_04A | chr16 | 71,016,992 | 4 | 83 | 4.8 |

| | | | | | |
|---|---|---|---|---|---|
| A7_16G | chr17 | 55,882,223 | 11 | 86 | 12.8 |
| A7_06C | chr18 | 10,699,389 | 5 | 66 | 7.6 |
| A7_06C | chr18 | 15,102,961 | 7 | 22 | 31.8 |
| A7_15T | chr18 | 34,460,701 | 2 | 36 | 5.6 |
| A7_15T | chr18 | 50,310,339 | 5 | 76 | 6.6 |
| A7_15A | chr18 | 63,110,753 | 7 | 70 | 10.0 |
| A7_15T | chr18 | 74,565,139 | 10 | 80 | 12.5 |
| A7_15T | chr19 | 46,759,432 | 9 | 59 | 15.3 |
| A7_06C | chr2 | 24,548,320 | 1 | 85 | 1.2 |
| A7_17A | chr2 | 24,987,998 | 18 | 114 | 15.8 |
| A7_16T | chr2 | 34,173,002 | 5 | 60 | 8.3 |
| A7_06A | chr2 | 52,479,453 | 5 | 40 | 12.5 |
| A7_15G | chr2 | 71,730,114 | 3 | 97 | 3.1 |
| A7_15G | chr2 | 81,146,960 | 6 | 100 | 6.0 |
| A7_13G | chr2 | 115,036,229 | 6 | 66 | 9.1 |
| A7_17C | chr2 | 117,954,822 | 7 | 61 | 11.5 |
| A7_05G | chr2 | 137,568,716 | 13 | 75 | 17.3 |
| A7_17A | chr20 | 5,369,013 | 13 | 102 | 12.7 |
| A7_16T | chr20 | 7,035,261 | 3 | 97 | 3.1 |
| A7_05G | chr20 | 52,220,406 | 3 | 76 | 3.9 |
| A7_No_Mismatch | chr23 | 5,399,999 | 0 | 44 | 0.0 |
| A7_No_Mismatch | chr23 | 34,335,321 | 2 | 34 | 5.9 |
| A7_16G | chr23 | 91,644,262 | 7 | 36 | 19.4 |
| A7_06A | chr23 | 94,195,717 | 1 | 32 | 3.1 |
| A7_15T | chr23 | 138,579,060 | 4 | 32 | 12.5 |
| A7_06A | chr23 | 151,085,876 | 2 | 49 | 4.1 |
| A7_15A | chr23 | 151,888,975 | 6 | 22 | 27.3 |
| A7_16G | chr24 | 4,883,524 | 7 | 36 | 19.4 |
| A7_05G | chr3 | 22,904,393 | 10 | 136 | 7.4 |
| A7_05C | chr3 | 35,464,128 | 3 | 62 | 4.8 |
| A7_04C | chr3 | 53,239,043 | 0 | 75 | 0.0 |
| A7_05G | chr3 | 55,354,158 | 5 | 70 | 7.1 |
| A7_05G | chr3 | 58,590,706 | 4 | 105 | 3.8 |
| A7_05G | chr3 | 78,134,639 | 7 | 59 | 11.9 |
| A7_15T | chr3 | 121,962,034 | 2 | 26 | 7.7 |
| A7_06C | chr4 | 7,604,262 | 2 | 70 | 2.9 |
| A7_No_Mismatch | chr4 | 66,626,203 | 2 | 42 | 4.8 |
| A7_06C | chr4 | 92,417,832 | 2 | 46 | 4.3 |
| A7_14G | chr4 | 107,705,340 | 7 | 56 | 12.5 |
| A7_05C | chr4 | 112,442,143 | 3 | 61 | 4.9 |
| A7_14G | chr4 | 133,577,266 | 3 | 42 | 7.1 |

| | | | | | |
|---|---|---|---|---|---|
| A7_14G | chr4 | 137,932,455 | 8 | 71 | 11.3 |
| A7_15A | chr4 | 150,057,469 | 6 | 67 | 9.0 |
| A7_14C | chr5 | 34,499,112 | 4 | 55 | 7.3 |
| A7_14C | chr5 | 43,697,842 | 7 | 83 | 8.4 |
| A7_14G | chr5 | 59,424,639 | 3 | 89 | 3.4 |
| A7_05C | chr5 | 67,394,178 | 1 | 52 | 1.9 |
| A7_05C | chr5 | 96,268,914 | 7 | 38 | 18.4 |
| A7_14G | chr5 | 111,764,536 | 5 | 84 | 6.0 |
| A7_06G | chr5 | 155,672,126 | 3 | 46 | 6.5 |
| A7_05C | chr5 | 180,342,021 | 2 | 55 | 3.6 |
| A7_14C | chr6 | 7,892,619 | 4 | 39 | 10.3 |
| A7_05G | chr6 | 98,141,170 | 3 | 49 | 6.1 |
| A7_05G | chr6 | 110,309,686 | 2 | 87 | 2.3 |
| A7_06G | chr6 | 158,768,156 | 7 | 68 | 10.3 |
| A7_05G | chr7 | 36,320,290 | 0 | 42 | 0.0 |
| A7_15A | chr7 | 38,114,597 | 6 | 80 | 7.5 |
| A7_13G | chr7 | 91,440,125 | 3 | 49 | 6.1 |
| A7_15G | chr7 | 124,548,627 | 5 | 90 | 5.6 |
| A7_No_Mismatch | chr7 | 133,713,677 | 0 | 69 | 0.0 |
| A7_05G | chr8 | 27,168,793 | 10 | 99 | 10.1 |
| A7_14G | chr8 | 28,630,172 | 4 | 77 | 5.2 |
| A7_16C | chr8 | 30,671,986 | 9 | 43 | 20.9 |
| A7_05G | chr8 | 102,934,894 | 1 | 47 | 2.1 |
| A7_14G | chr8 | 104,102,765 | 5 | 71 | 7.0 |
| A7_15G | chr8 | 114,113,528 | 4 | 41 | 9.8 |
| A7_05G | chr8 | 123,672,981 | 11 | 82 | 13.4 |
| A7_06G | chr9 | 78,290,171 | 7 | 71 | 9.9 |
| A7_05G | chr9 | 138,173,180 | 14 | 84 | 16.7 |
