## Supplemental length data for "CRISPR/Cas9 Optimization Using Electronic Genome Mapping: Potential Role for Studying Human Genetic Disease"

| gRNA | Chromosome | hg38 Position | tags | Molecules | % Tagged | Distance |
|---|---|---|---|---|---|---|
| A1 | chr18 | 5,270,508 | 15 | 81 | 18.5 | 4 |
| B5 | chr14 | 56,425,539 | 8 | 32 | 25.0 | 4 |
| B5 | chr20 | 43,002,084 | 20 | 90 | 27.8 | 4 |
| B5 | chr11 | 101,708,756 | 32 | 41 | 78.0 | 4 |
| B3_HK5671 | chr14 | 66,731,008 | 3 | 52 | 5.8 | 5 |
| B3_AU1054 | chr7 | 39,633,474 | 4 | 66 | 6.1 | 5 |
| B3_HK5671 | chr7 | 39,633,474 | 7 | 55 | 12.7 | 5 |
| B3_AU1054 | chr14 | 89,219,233 | 15 | 102 | 14.7 | 5 |
| B3_AU1054 | chr6 | 135,079,313 | 13 | 87 | 14.9 | 5 |
| B3_HK5671 | chr2 | 170,968,780 | 13 | 81 | 16.0 | 5 |
| B3_AU1054 | chr2 | 170,968,780 | 16 | 97 | 16.5 | 5 |
| B3_AU1054 | chr13 | 61,354,596 | 9 | 49 | 18.4 | 5 |
| B3_AU1054 | chr14 | 66,731,008 | 8 | 39 | 20.5 | 5 |
| B3_HK5671 | chr14 | 89,219,233 | 21 | 98 | 21.4 | 5 |
| B3_HK5671 | chr6 | 135,079,313 | 13 | 49 | 26.5 | 5 |
| B3_HK5671 | chr2 | 20,427,360 | 27 | 88 | 30.7 | 5 |
| B3_HK5671 | chr4 | 90,848,820 | 23 | 66 | 34.8 | 5 |
| B3_HK5671 | chr6 | 120,005,689 | 16 | 45 | 35.6 | 5 |
| B3_AU1054 | chr2 | 20,427,360 | 54 | 144 | 37.5 | 5 |
| B3_HK5671 | chr12 | 91,784,782 | 26 | 68 | 38.2 | 5 |
| A7 | chr18 | 5,270,509 | 29 | 71 | 40.8 | 5 |
| B3_HK5671 | chr6 | 153,563,127 | 20 | 48 | 41.7 | 5 |
| B3_AU1054 | chr8 | 130,512,592 | 28 | 64 | 43.8 | 5 |
| B3_HK5671 | chr23 | 147,875,283 | 16 | 36 | 44.4 | 5 |
| B3_AU1054 | chr4 | 90,848,820 | 37 | 72 | 51.4 | 5 |
| B3_AU1054 | chr6 | 120,005,689 | 31 | 56 | 55.4 | 5 |
| B3_HK5671 | chr13 | 25,065,643 | 49 | 75 | 65.3 | 5 |
| B3_AU1054 | chr6 | 153,563,127 | 38 | 57 | 66.7 | 5 |
| B3_HK5671 | chr17 | 4,340,550 | 34 | 51 | 66.7 | 5 |
| B3_HK5671 | chr13 | 61,354,596 | 33 | 49 | 67.3 | 5 |
| B3_HK5671 | chr2 | 82,755,723 | 36 | 52 | 69.2 | 5 |
| B3_AU1054 | chr12 | 91,784,782 | 50 | 71 | 70.4 | 5 |
| B3_HK5671 | chr8 | 130,512,592 | 59 | 82 | 72.0 | 5 |
| B3_AU1054 | chr13 | 25,065,643 | 67 | 89 | 75.3 | 5 |
| B3_AU1054 | chr17 | 4,340,550 | 48 | 63 | 76.2 | 5 |
| B3_AU1054 | chr2 | 82,755,723 | 45 | 59 | 76.3 | 5 |
| B3_AU1054 | chr14 | 33,217,158 | 59 | 76 | 77.6 | 5 |
| B3_AU1054 | chr10 | 120,191,715 | 43 | 54 | 79.6 | 5 |
| B3_AU1054 | chr23 | 147,875,283 | 37 | 45 | 82.2 | 5 |
| B3_HK5671 | chr14 | 33,217,158 | 80 | 90 | 88.9 | 5 |

| | | | | | | |
|---|---|---|---|---|---|---|
| B3_HK5671 | chr10 | 120,191,715 | 39 | 43 | 90.7 | 5 |
| B3_HK5671 | chr2 | 8,344,008 | 90 | 96 | 93.8 | 5 |
| B3_AU1054 | chr2 | 8,344,008 | 103 | 107 | 96.3 | 5 |
| A1 | chr14 | 66,730,964 | 3 | 66 | 4.5 | 6 |
| A1 | chr13 | 61,354,552 | 3 | 59 | 5.1 | 6 |
| A1 | chr17 | 4,340,506 | 5 | 69 | 7.2 | 6 |
| A1 | chr2 | 82,755,744 | 5 | 54 | 9.3 | 6 |
| A1 | chr6 | 110,607,014 | 7 | 64 | 10.9 | 6 |
| A1 | chr9 | 92,127,061 | 11 | 91 | 12.1 | 6 |
| A1 | chr2 | 176,265,352 | 8 | 65 | 12.3 | 6 |
| A1 | chr17 | 14,468,251 | 10 | 80 | 12.5 | 6 |
| A1 | chr8 | 41,425,274 | 10 | 72 | 13.9 | 6 |
| A1 | chr10 | 26,001,046 | 9 | 57 | 15.8 | 6 |
| A1 | chr7 | 94,175,018 | 12 | 76 | 15.8 | 6 |
| A5 | chr14 | 56,425,519 | 16 | 74 | 21.6 | 6 |
| A1 | chr4 | 18,053,105 | 13 | 58 | 22.4 | 6 |
| A1 | chr11 | 115,652,641 | 19 | 72 | 26.4 | 6 |
| A7 | chr9 | 138,033,806 | 48 | 71 | 67.6 | 6 |
| A2 | chr9 | 92,202,584 | 70 | 96 | 72.9 | 6 |
| A7 | chr3 | 116,653,046 | 25 | 34 | 73.5 | 6 |
| A2 | chr6 | 84,300,717 | 48 | 62 | 77.4 | 6 |
| A7 | chr11 | 101,708,713 | 47 | 53 | 88.7 | 6 |
| A1 | chr6 | 32,985,852 | 83 | 84 | 98.8 | 6 |
| A7 | chr17 | 4,340,506 | 5 | 57 | 8.8 | 7 |
| A7 | chr14 | 66,730,964 | 6 | 47 | 12.8 | 7 |
| A7 | chr7 | 94,175,018 | 9 | 50 | 18.0 | 7 |
| A7 | chr6 | 135,079,269 | 15 | 53 | 28.3 | 7 |
| A7 | chr2 | 8,344,052 | 25 | 86 | 29.1 | 7 |
| A7 | chr8 | 41,425,274 | 25 | 70 | 35.7 | 7 |
| A7 | chr9 | 122,634,606 | 26 | 63 | 41.3 | 7 |
| A7 | chr12 | 114,623,246 | 45 | 67 | 67.2 | 7 |
| A7 | chr13 | 61,354,552 | 41 | 53 | 77.4 | 7 |
| A7 | chr11 | 115,652,641 | 57 | 73 | 78.1 | 7 |
| A5 | chr8 | 130,512,550 | 47 | 59 | 79.7 | 7 |
| A7 | chr2 | 20,427,316 | 76 | 95 | 80.0 | 7 |
| A7 | chr23 | 46,286,328 | 47 | 56 | 83.9 | 7 |
| A7 | chr4 | 18,053,105 | 53 | 58 | 91.4 | 7 |
| A7 | chr6 | 32,985,853 | 58 | 59 | 98.3 | 7 |
| B5 | chr23 | 96,465,891 | 16 | 25 | 64.0 | 8 |
| B5 | chr2 | 238,051,501 | 42 | 58 | 72.4 | 8 |
| B5 | chr21 | 45,719,605 | 31 | 39 | 79.5 | 8 |

| | | | | | | |
|---|---|---|---|---|---|---|
| B5 | chr14 | 103,813,517 | 42 | 49 | 85.7 | 8 |
| B5 | chr15 | 25,856,280 | 45 | 51 | 88.2 | 8 |
| A1 | chr13 | 46,840,994 | 66 | 72 | 91.7 | 8 |
| B5 | chr9 | 16,983,805 | 50 | 54 | 92.6 | 8 |
| B5 | chr8 | 40,550,646 | 36 | 38 | 94.7 | 8 |
| B5 | chr8 | 35,321,431 | 35 | 35 | 100.0 | 8 |
| A1 | chr2 | 87,218,288 | 11 | 66 | 16.7 | 9 |
| B3_AU1054 | chr10 | 28,727,511 | 21 | 99 | 21.2 | 9 |
| B3_HK5671 | chr10 | 28,727,511 | 61 | 94 | 64.9 | 9 |
| B3_AU1054 | chr3 | 12,784,234 | 93 | 136 | 68.4 | 9 |
| A1 | chr6 | 169,560,169 | 35 | 51 | 68.6 | 9 |
| B3_HK5671 | chr3 | 12,784,234 | 72 | 103 | 69.9 | 9 |
| B3_HK5671 | chr10 | 22,218,046 | 72 | 98 | 73.5 | 9 |
| A1 | chr23 | 151,149,027 | 26 | 34 | 76.5 | 9 |
| A1 | chr13 | 49,475,219 | 45 | 58 | 77.6 | 9 |
| A1 | chr6 | 126,408,450 | 43 | 55 | 78.2 | 9 |
| A1 | chr3 | 191,425,447 | 55 | 68 | 80.9 | 9 |
| A1 | chr6 | 167,981,980 | 43 | 53 | 81.1 | 9 |
| A1 | chr8 | 78,558,582 | 35 | 43 | 81.4 | 9 |
| B3_AU1054 | chr10 | 22,218,046 | 76 | 93 | 81.7 | 9 |
| A1 | chr23 | 32,757,553 | 44 | 52 | 84.6 | 9 |
| A1 | chr7 | 124,086,100 | 51 | 60 | 85.0 | 9 |
| A1 | chr21 | 45,719,558 | 60 | 70 | 85.7 | 9 |
| A1 | chr15 | 88,314,760 | 73 | 85 | 85.9 | 9 |
| A1 | chr1 | 100,779,922 | 50 | 58 | 86.2 | 9 |
| A1 | chr2 | 188,009,502 | 58 | 67 | 86.6 | 9 |
| A1 | chr7 | 85,043,688 | 47 | 54 | 87.0 | 9 |
| A1 | chr5 | 35,729,778 | 48 | 55 | 87.3 | 9 |
| A1 | chr11 | 39,262,712 | 32 | 36 | 88.9 | 9 |
| A1 | chr5 | 159,219,798 | 48 | 54 | 88.9 | 9 |
| A1 | chr7 | 151,881,493 | 48 | 54 | 88.9 | 9 |
| B3_HK5671 | chr7 | 37,273,367 | 57 | 64 | 89.1 | 9 |
| A1 | chr23 | 96,465,937 | 25 | 28 | 89.3 | 9 |
| A1 | chr9 | 31,578,527 | 51 | 57 | 89.5 | 9 |
| B3_AU1054 | chr7 | 37,273,367 | 43 | 48 | 89.6 | 9 |
| B3_AU1054 | chr3 | 156,999,035 | 96 | 107 | 89.7 | 9 |
| A1 | chr4 | 117,139,033 | 53 | 59 | 89.8 | 9 |
| B3_AU1054 | chr5 | 132,525,048 | 71 | 78 | 91.0 | 9 |
| B3_AU1054 | chr5 | 81,407,970 | 65 | 71 | 91.5 | 9 |
| B3_HK5671 | chr14 | 85,688,679 | 43 | 47 | 91.5 | 9 |
| A1 | chr2 | 214,016,394 | 11 | 12 | 91.7 | 9 |

| | | | | | | |
|---|---|---|---|---|---|---|
| A1 | chr7 | 69,499,590 | 58 | 63 | 92.1 | 9 |
| A7 | chr13 | 46,840,994 | 58 | 63 | 92.1 | 9 |
| A1 | chr14 | 94,268,795 | 59 | 64 | 92.2 | 9 |
| B3_AU1054 | chr19 | 33,923,757 | 72 | 78 | 92.3 | 9 |
| B3_HK5671 | chr4 | 160,855,842 | 36 | 39 | 92.3 | 9 |
| A1 | chr8 | 90,890,237 | 60 | 65 | 92.3 | 9 |
| B3_HK5671 | chr2 | 174,921,770 | 88 | 95 | 92.6 | 9 |
| B3_HK5671 | chr12 | 92,654,467 | 92 | 99 | 92.9 | 9 |
| A1 | chr21 | 15,317,160 | 58 | 62 | 93.5 | 9 |
| B3_AU1054 | chr7 | 94,249,754 | 59 | 63 | 93.7 | 9 |
| B3_HK5671 | chr23 | 36,690,465 | 59 | 63 | 93.7 | 9 |
| B3_HK5671 | chr6 | 4,678,721 | 61 | 65 | 93.8 | 9 |
| A1 | chr9 | 16,983,758 | 63 | 67 | 94.0 | 9 |
| B3_HK5671 | chr5 | 81,407,970 | 65 | 69 | 94.2 | 9 |
| A1 | chr9 | 135,911,458 | 49 | 52 | 94.2 | 9 |
| B3_AU1054 | chr10 | 127,591,710 | 100 | 106 | 94.3 | 9 |
| B3_AU1054 | chr6 | 4,678,721 | 83 | 88 | 94.3 | 9 |
| B3_AU1054 | chr4 | 160,855,842 | 51 | 54 | 94.4 | 9 |
| B3_AU1054 | chr5 | 142,182,017 | 68 | 72 | 94.4 | 9 |
| B3_AU1054 | chr2 | 196,108,283 | 69 | 73 | 94.5 | 9 |
| B3_AU1054 | chr1 | 18,579,475 | 53 | 56 | 94.6 | 9 |
| B3_AU1054 | chr13 | 50,329,589 | 71 | 75 | 94.7 | 9 |
| B3_HK5671 | chr3 | 31,314,738 | 93 | 98 | 94.9 | 9 |
| B3_HK5671 | chr5 | 84,732,889 | 37 | 39 | 94.9 | 9 |
| B3_HK5671 | chr15 | 75,755,050 | 76 | 80 | 95.0 | 9 |
| B3_AU1054 | chr23 | 71,649,227 | 79 | 83 | 95.2 | 9 |
| A1 | chr2 | 53,194,591 | 60 | 63 | 95.2 | 9 |
| B3_HK5671 | chr17 | 62,884,910 | 61 | 64 | 95.3 | 9 |
| A1 | chr14 | 62,513,255 | 63 | 66 | 95.5 | 9 |
| A1 | chr23 | 5,695,485 | 43 | 45 | 95.6 | 9 |
| B3_HK5671 | chr2 | 196,108,283 | 65 | 68 | 95.6 | 9 |
| B3_AU1054 | chr14 | 85,688,679 | 46 | 48 | 95.8 | 9 |
| B3_HK5671 | chr4 | 7,977,145 | 50 | 52 | 96.2 | 9 |
| B3_AU1054 | chr14 | 35,925,533 | 52 | 54 | 96.3 | 9 |
| A1 | chr19 | 33,614,108 | 80 | 83 | 96.4 | 9 |
| B3_HK5671 | chr7 | 94,249,754 | 54 | 56 | 96.4 | 9 |
| B3_AU1054 | chr3 | 79,740,608 | 110 | 114 | 96.5 | 9 |
| B3_HK5671 | chr14 | 35,925,533 | 55 | 57 | 96.5 | 9 |
| B3_HK5671 | chr8 | 15,966,109 | 82 | 85 | 96.5 | 9 |
| B3_AU1054 | chr18 | 59,993,315 | 86 | 89 | 96.6 | 9 |
| B3_HK5671 | chr19 | 33,923,757 | 56 | 58 | 96.6 | 9 |

| | | | | | | |
|---|---|---|---|---|---|---|
| B3_HK5671 | chr1 | 18,579,475 | 59 | 61 | 96.7 | 9 |
| B3_HK5671 | chr17 | 72,979,152 | 58 | 60 | 96.7 | 9 |
| B3_AU1054 | chr17 | 62,884,910 | 60 | 62 | 96.8 | 9 |
| B3_AU1054 | chr17 | 72,979,152 | 91 | 94 | 96.8 | 9 |
| B3_AU1054 | chr5 | 84,732,889 | 60 | 62 | 96.8 | 9 |
| B3_HK5671 | chr1 | 85,373,510 | 61 | 63 | 96.8 | 9 |
| B3_HK5671 | chr2 | 112,457,935 | 60 | 62 | 96.8 | 9 |
| B3_AU1054 | chr1 | 85,373,510 | 95 | 98 | 96.9 | 9 |
| B3_AU1054 | chr8 | 15,966,109 | 94 | 97 | 96.9 | 9 |
| B3_AU1054 | chr3 | 31,314,738 | 96 | 99 | 97.0 | 9 |
| B3_AU1054 | chr16 | 72,174,748 | 100 | 103 | 97.1 | 9 |
| B3_HK5671 | chr20 | 8,374,235 | 67 | 69 | 97.1 | 9 |
| B3_AU1054 | chr22 | 35,231,088 | 35 | 36 | 97.2 | 9 |
| B3_AU1054 | chr3 | 78,389,253 | 69 | 71 | 97.2 | 9 |
| A1 | chr3 | 189,196,603 | 70 | 72 | 97.2 | 9 |
| B3_HK5671 | chr22 | 35,231,088 | 36 | 37 | 97.3 | 9 |
| B3_AU1054 | chr15 | 75,755,050 | 74 | 76 | 97.4 | 9 |
| B3_AU1054 | chr17 | 34,471,434 | 83 | 85 | 97.6 | 9 |
| A1 | chr6 | 71,663,044 | 41 | 42 | 97.6 | 9 |
| A1 | chr16 | 78,933,771 | 83 | 85 | 97.6 | 9 |
| A1 | chr2 | 167,162,547 | 83 | 85 | 97.6 | 9 |
| B3_HK5671 | chr3 | 156,999,035 | 84 | 86 | 97.7 | 9 |
| B3_AU1054 | chr3 | 82,866,578 | 45 | 46 | 97.8 | 9 |
| B3_HK5671 | chr16 | 72,174,748 | 89 | 91 | 97.8 | 9 |
| B3_AU1054 | chr2 | 174,921,770 | 98 | 100 | 98.0 | 9 |
| B3_HK5671 | chr5 | 132,525,048 | 97 | 99 | 98.0 | 9 |
| B3_AU1054 | chr12 | 92,654,467 | 103 | 105 | 98.1 | 9 |
| B3_AU1054 | chr2 | 112,457,935 | 52 | 53 | 98.1 | 9 |
| B3_HK5671 | chr10 | 127,591,710 | 52 | 53 | 98.1 | 9 |
| B3_AU1054 | chr6 | 117,282,416 | 56 | 57 | 98.2 | 9 |
| B3_AU1054 | chr23 | 36,690,465 | 59 | 60 | 98.3 | 9 |
| A1 | chr3 | 3,317,605 | 61 | 62 | 98.4 | 9 |
| B3_AU1054 | chr20 | 8,374,235 | 62 | 63 | 98.4 | 9 |
| B3_HK5671 | chr5 | 142,182,017 | 62 | 63 | 98.4 | 9 |
| A1 | chr5 | 171,977,807 | 62 | 63 | 98.4 | 9 |
| A1 | chr9 | 122,458,644 | 64 | 65 | 98.5 | 9 |
| B3_AU1054 | chr2 | 150,585,242 | 65 | 66 | 98.5 | 9 |
| B3_HK5671 | chr3 | 78,389,253 | 64 | 65 | 98.5 | 9 |
| B3_HK5671 | chr2 | 150,585,242 | 70 | 71 | 98.6 | 9 |
| B3_AU1054 | chr18 | 24,107,546 | 74 | 75 | 98.7 | 9 |
| B3_AU1054 | chr20 | 44,743,452 | 78 | 79 | 98.7 | 9 |

| | | | | | | |
|---|---|---|---|---|---|---|
| B3_AU1054 | chr8 | 131,021,677 | 74 | 75 | 98.7 | 9 |
| B3_HK5671 | chr13 | 50,329,589 | 74 | 75 | 98.7 | 9 |
| B3_HK5671 | chr3 | 79,740,608 | 77 | 78 | 98.7 | 9 |
| B3_AU1054 | chr4 | 7,977,145 | 80 | 81 | 98.8 | 9 |
| B3_HK5671 | chr18 | 24,107,546 | 85 | 86 | 98.8 | 9 |
| B3_HK5671 | chr18 | 59,993,315 | 83 | 84 | 98.8 | 9 |
| B3_HK5671 | chr7 | 26,501,327 | 88 | 89 | 98.9 | 9 |
| B3_AU1054 | chr7 | 26,501,327 | 113 | 114 | 99.1 | 9 |
| A1 | chr23 | 138,032,813 | 19 | 19 | 100.0 | 9 |
| A1 | chr4 | 62,382,542 | 12 | 12 | 100.0 | 9 |
| B3_AU1054 | chr5 | 170,997,103 | 48 | 48 | 100.0 | 9 |
| B3_HK5671 | chr17 | 34,471,434 | 58 | 58 | 100.0 | 9 |
| B3_HK5671 | chr20 | 44,743,452 | 66 | 66 | 100.0 | 9 |
| B3_HK5671 | chr23 | 71,649,227 | 50 | 50 | 100.0 | 9 |
| B3_HK5671 | chr3 | 82,866,578 | 15 | 15 | 100.0 | 9 |
| B3_HK5671 | chr5 | 170,997,103 | 32 | 32 | 100.0 | 9 |
| B3_HK5671 | chr6 | 117,282,416 | 51 | 51 | 100.0 | 9 |
| B3_HK5671 | chr8 | 25,358,319 | 60 | 60 | 100.0 | 9 |
| B3_HK5671 | chr8 | 131,021,677 | 64 | 64 | 100.0 | 9 |
| A7 | chr5 | 53,387,386 | 12 | 51 | 23.5 | 10 |
| A2 | chr12 | 24,575,103 | 16 | 63 | 25.4 | 10 |
| A7 | chr17 | 74,171,602 | 31 | 87 | 35.6 | 10 |
| A5 | chr7 | 156,537,473 | 64 | 103 | 62.1 | 10 |
| A7 | chr6 | 167,981,980 | 53 | 74 | 71.6 | 10 |
| A5 | chr13 | 97,621,220 | 64 | 85 | 75.3 | 10 |
| A5 | chr20 | 53,429,435 | 68 | 89 | 76.4 | 10 |
| A5 | chr20 | 18,627,806 | 59 | 77 | 76.6 | 10 |
| A1 | chr7 | 106,423,041 | 53 | 66 | 80.3 | 10 |
| A7 | chr21 | 45,719,558 | 49 | 60 | 81.7 | 10 |
| A7 | chr5 | 159,219,798 | 46 | 56 | 82.1 | 10 |
| A5 | chr1 | 214,203,971 | 49 | 59 | 83.1 | 10 |
| A5 | chr6 | 78,598,035 | 60 | 72 | 83.3 | 10 |
| B3_HK5671 | chr14 | 56,344,938 | 56 | 67 | 83.6 | 10 |
| B3_AU1054 | chr14 | 56,344,938 | 76 | 90 | 84.4 | 10 |
| A5 | chr1 | 99,482,845 | 51 | 60 | 85.0 | 10 |
| A1 | chr1 | 61,346,982 | 57 | 67 | 85.1 | 10 |
| A7 | chr9 | 16,983,758 | 63 | 74 | 85.1 | 10 |
| A1 | chr3 | 186,002,693 | 58 | 68 | 85.3 | 10 |
| A5 | chr3 | 14,087,719 | 71 | 83 | 85.5 | 10 |
| A5 | chr7 | 139,977,386 | 98 | 114 | 86.0 | 10 |
| A5 | chr21 | 44,776,666 | 39 | 45 | 86.7 | 10 |

| | | | | | | |
|---|---|---|---|---|---|---|
| A7 | chr23 | 152,562,747 | 20 | 23 | 87.0 | 10 |
| A1 | chr16 | 62,499,265 | 47 | 54 | 87.0 | 10 |
| A7 | chr1 | 100,779,923 | 57 | 65 | 87.7 | 10 |
| A7 | chr7 | 69,499,590 | 75 | 85 | 88.2 | 10 |
| A7 | chr7 | 85,043,688 | 39 | 44 | 88.6 | 10 |
| A5 | chr12 | 80,777,415 | 55 | 62 | 88.7 | 10 |
| A7 | chr14 | 94,268,796 | 56 | 63 | 88.9 | 10 |
| A5 | chr4 | 33,795,506 | 81 | 91 | 89.0 | 10 |
| A5 | chr14 | 24,604,918 | 74 | 83 | 89.2 | 10 |
| A2 | chr23 | 6,373,683 | 17 | 19 | 89.5 | 10 |
| A1 | chr3 | 12,258,548 | 60 | 67 | 89.6 | 10 |
| A7 | chr6 | 24,754,689 | 61 | 68 | 89.7 | 10 |
| A5 | chr16 | 77,631,963 | 70 | 78 | 89.7 | 10 |
| A5 | chr2 | 55,091,906 | 93 | 103 | 90.3 | 10 |
| B3_HK5671 | chr21 | 31,767,146 | 48 | 53 | 90.6 | 10 |
| A7 | chr6 | 126,408,451 | 40 | 44 | 90.9 | 10 |
| A7 | chr2 | 167,162,548 | 83 | 91 | 91.2 | 10 |
| A7 | chr9 | 135,911,459 | 44 | 48 | 91.7 | 10 |
| A7 | chr16 | 16,478,250 | 92 | 100 | 92.0 | 10 |
| A1 | chr23 | 13,450,913 | 24 | 26 | 92.3 | 10 |
| A7 | chr7 | 151,881,494 | 63 | 68 | 92.6 | 10 |
| A1 | chr1 | 106,542,707 | 41 | 44 | 93.2 | 10 |
| A5 | chr7 | 156,370,004 | 55 | 59 | 93.2 | 10 |
| A7 | chr5 | 171,977,808 | 58 | 62 | 93.5 | 10 |
| A5 | chr8 | 37,580,756 | 122 | 130 | 93.8 | 10 |
| A7 | chr16 | 78,933,772 | 92 | 98 | 93.9 | 10 |
| B3_AU1054 | chr21 | 31,767,146 | 77 | 82 | 93.9 | 10 |
| A1 | chr4 | 111,802,808 | 48 | 51 | 94.1 | 10 |
| A1 | chr9 | 118,913,308 | 50 | 53 | 94.3 | 10 |
| A1 | chr15 | 52,003,266 | 51 | 54 | 94.4 | 10 |
| A7 | chr2 | 214,016,395 | 17 | 18 | 94.4 | 10 |
| A1 | chr7 | 94,249,801 | 71 | 75 | 94.7 | 10 |
| A7 | chr21 | 15,317,160 | 71 | 75 | 94.7 | 10 |
| A7 | chr23 | 32,757,553 | 56 | 59 | 94.9 | 10 |
| A7 | chr12 | 63,131,437 | 38 | 40 | 95.0 | 10 |
| A7 | chr2 | 75,542,038 | 97 | 102 | 95.1 | 10 |
| A1 | chr10 | 105,970,232 | 59 | 62 | 95.2 | 10 |
| A1 | chr18 | 59,993,267 | 83 | 87 | 95.4 | 10 |
| A1 | chr10 | 49,791,271 | 63 | 66 | 95.5 | 10 |
| A5 | chr8 | 40,927,530 | 43 | 45 | 95.6 | 10 |
| A7 | chr19 | 33,614,108 | 109 | 114 | 95.6 | 10 |

| | | | | | | |
|---|---|---|---|---|---|---|
| B3_HK5671 | chr7 | 154,672,145 | 45 | 47 | 95.7 | 10 |
| A1 | chr13 | 96,621,313 | 45 | 47 | 95.7 | 10 |
| B3_AU1054 | chr10 | 92,168,638 | 68 | 71 | 95.8 | 10 |
| A1 | chr1 | 227,387,479 | 70 | 73 | 95.9 | 10 |
| A1 | chr6 | 24,589,520 | 71 | 74 | 95.9 | 10 |
| A2 | chr5 | 10,200,588 | 49 | 51 | 96.1 | 10 |
| A1 | chr8 | 96,454,053 | 75 | 78 | 96.2 | 10 |
| A7 | chr9 | 122,458,645 | 56 | 58 | 96.6 | 10 |
| A5 | chr13 | 53,247,656 | 94 | 97 | 96.9 | 10 |
| A1 | chr23 | 5,863,190 | 34 | 35 | 97.1 | 10 |
| A2 | chr16 | 51,627,000 | 70 | 72 | 97.2 | 10 |
| A1 | chr17 | 34,471,481 | 72 | 74 | 97.3 | 10 |
| A7 | chr2 | 53,194,592 | 77 | 79 | 97.5 | 10 |
| A1 | chr13 | 90,556,610 | 78 | 80 | 97.5 | 10 |
| A5 | chr2 | 111,747,533 | 49 | 50 | 98.0 | 10 |
| A7 | chr2 | 23,304,137 | 99 | 101 | 98.0 | 10 |
| A5 | chr2 | 139,219,409 | 57 | 58 | 98.3 | 10 |
| A1 | chr3 | 82,866,588 | 59 | 60 | 98.3 | 10 |
| B3_AU1054 | chr7 | 154,672,145 | 72 | 73 | 98.6 | 10 |
| A2 | chr1 | 14,656,771 | 82 | 83 | 98.8 | 10 |
| A1 | chr16 | 54,874,242 | 77 | 77 | 100.0 | 10 |
| A1 | chr5 | 170,997,055 | 39 | 39 | 100.0 | 10 |
| A7 | chr10 | 7,271,890 | 16 | 16 | 100.0 | 10 |
| A7 | chr23 | 151,149,028 | 36 | 36 | 100.0 | 10 |
| A7 | chr11 | 39,262,713 | 61 | 61 | 100.0 | 10 |
| B3_HK5671 | chr10 | 92,168,638 | 57 | 57 | 100.0 | 10 |
| A5 | chr4 | 112,112,326 | 28 | 67 | 41.8 | 11 |
| A7 | chr21 | 40,591,057 | 47 | 64 | 73.4 | 11 |
| A7 | chr12 | 69,424,622 | 40 | 54 | 74.1 | 11 |
| A7 | chr4 | 11,032,345 | 60 | 78 | 76.9 | 11 |
| A7 | chr15 | 92,015,487 | 48 | 62 | 77.4 | 11 |
| A7 | chr7 | 76,646,778 | 54 | 68 | 79.4 | 11 |
| A5 | chr2 | 186,443,210 | 41 | 51 | 80.4 | 11 |
| A7 | chr16 | 72,174,796 | 61 | 75 | 81.3 | 11 |
| A7 | chr7 | 106,423,042 | 54 | 65 | 83.1 | 11 |
| A7 | chr18 | 59,993,267 | 69 | 82 | 84.1 | 11 |
| A5 | chr2 | 8,838,904 | 74 | 87 | 85.1 | 11 |
| A7 | chr15 | 52,003,267 | 40 | 47 | 85.1 | 11 |
| A7 | chr17 | 72,979,104 | 64 | 75 | 85.3 | 11 |
| A7 | chr13 | 55,782,847 | 43 | 50 | 86.0 | 11 |
| B3_AU1054 | chr12 | 115,783,841 | 71 | 82 | 86.6 | 11 |

| | | | | | | |
|---|---|---|---|---|---|---|
| A7 | chr8 | 96,454,054 | 65 | 74 | 87.8 | 11 |
| A7 | chr3 | 186,002,693 | 75 | 85 | 88.2 | 11 |
| A7 | chr2 | 19,892,219 | 40 | 45 | 88.9 | 11 |
| A7 | chr12 | 106,513,396 | 34 | 38 | 89.5 | 11 |
| A7 | chr10 | 28,727,463 | 92 | 102 | 90.2 | 11 |
| A7 | chr1 | 61,346,983 | 48 | 53 | 90.6 | 11 |
| A5 | chr15 | 78,747,901 | 53 | 58 | 91.4 | 11 |
| A7 | chr16 | 62,499,265 | 40 | 43 | 93.0 | 11 |
| A7 | chr3 | 82,866,626 | 41 | 44 | 93.2 | 11 |
| A7 | chr4 | 64,768,595 | 43 | 46 | 93.5 | 11 |
| A7 | chr7 | 94,249,802 | 67 | 71 | 94.4 | 11 |
| A7 | chr11 | 45,056,130 | 38 | 40 | 95.0 | 11 |
| A7 | chr13 | 96,621,313 | 60 | 63 | 95.2 | 11 |
| A7 | chr5 | 170,997,055 | 21 | 22 | 95.5 | 11 |
| A7 | chr6 | 24,589,520 | 66 | 69 | 95.7 | 11 |
| A5 | chr15 | 77,948,100 | 47 | 49 | 95.9 | 11 |
| A7 | chr7 | 113,173,651 | 89 | 92 | 96.7 | 11 |
| A7 | chr17 | 34,471,482 | 45 | 46 | 97.8 | 11 |
| A7 | chr13 | 36,603,278 | 49 | 50 | 98.0 | 11 |
| A7 | chr11 | 29,983,191 | 64 | 65 | 98.5 | 11 |
| B3_HK5671 | chr12 | 115,783,841 | 55 | 55 | 100.0 | 11 |
| B3_HK5671 | chr3 | 142,267,054 | 61 | 74 | 82.4 | 12 |
| B3_AU1054 | chr3 | 142,267,054 | 72 | 85 | 84.7 | 12 |
| B3_HK5671 | chr2 | 211,100,457 | 17 | 20 | 85.0 | 12 |
| B3_AU1054 | chr16 | 66,227,809 | 62 | 71 | 87.3 | 12 |
| B3_AU1054 | chr9 | 113,704,117 | 106 | 120 | 88.3 | 12 |
| B3_AU1054 | chr8 | 122,860,180 | 78 | 88 | 88.6 | 12 |
| B3_HK5671 | chr7 | 67,085,382 | 87 | 95 | 91.6 | 12 |
| B3_AU1054 | chr2 | 211,100,457 | 23 | 25 | 92.0 | 12 |
| B3_AU1054 | chr14 | 20,358,548 | 63 | 68 | 92.6 | 12 |
| B3_AU1054 | chr6 | 22,130,931 | 87 | 94 | 92.6 | 12 |
| B3_AU1054 | chr6 | 163,225,362 | 103 | 111 | 92.8 | 12 |
| B3_AU1054 | chr13 | 78,117,061 | 54 | 58 | 93.1 | 12 |
| B3_HK5671 | chr6 | 103,796,054 | 54 | 58 | 93.1 | 12 |
| B3_AU1054 | chr12 | 22,106,159 | 42 | 45 | 93.3 | 12 |
| B3_AU1054 | chr6 | 46,257,577 | 97 | 104 | 93.3 | 12 |
| B3_HK5671 | chr12 | 105,657,586 | 112 | 120 | 93.3 | 12 |
| B3_HK5671 | chr9 | 113,704,117 | 85 | 91 | 93.4 | 12 |
| B3_AU1054 | chr23 | 136,704,669 | 80 | 85 | 94.1 | 12 |
| B3_AU1054 | chr2 | 124,330,527 | 81 | 86 | 94.2 | 12 |
| B3_AU1054 | chr12 | 105,657,586 | 132 | 140 | 94.3 | 12 |

| | | | | | | |
|---|---|---|---|---|---|---|
| B3_AU1054 | chr6 | 103,796,054 | 66 | 70 | 94.3 | 12 |
| B3_HK5671 | chr16 | 46,481,122 | 67 | 71 | 94.4 | 12 |
| B3_AU1054 | chr13 | 48,590,627 | 89 | 94 | 94.7 | 12 |
| B3_AU1054 | chr5 | 126,036,055 | 94 | 99 | 94.9 | 12 |
| B3_AU1054 | chr2 | 143,534,188 | 83 | 87 | 95.4 | 12 |
| B3_HK5671 | chr13 | 48,590,627 | 64 | 67 | 95.5 | 12 |
| B3_HK5671 | chr16 | 66,227,809 | 84 | 88 | 95.5 | 12 |
| B3_AU1054 | chr5 | 128,659,574 | 65 | 68 | 95.6 | 12 |
| B3_AU1054 | chr13 | 74,453,180 | 45 | 47 | 95.7 | 12 |
| B3_AU1054 | chr5 | 166,150,005 | 76 | 79 | 96.2 | 12 |
| B3_AU1054 | chr7 | 67,085,382 | 75 | 78 | 96.2 | 12 |
| B3_HK5671 | chr12 | 22,106,159 | 75 | 78 | 96.2 | 12 |
| B3_HK5671 | chr2 | 124,330,527 | 53 | 55 | 96.4 | 12 |
| B3_HK5671 | chr1 | 87,994,642 | 56 | 58 | 96.6 | 12 |
| B3_AU1054 | chr21 | 35,303,564 | 88 | 91 | 96.7 | 12 |
| B3_HK5671 | chr14 | 20,358,548 | 61 | 63 | 96.8 | 12 |
| B3_HK5671 | chr5 | 126,036,055 | 91 | 94 | 96.8 | 12 |
| B3_HK5671 | chr6 | 22,130,931 | 90 | 93 | 96.8 | 12 |
| B3_HK5671 | chr13 | 78,117,061 | 34 | 35 | 97.1 | 12 |
| B3_HK5671 | chr21 | 35,303,564 | 102 | 105 | 97.1 | 12 |
| B3_AU1054 | chr12 | 67,968,041 | 69 | 71 | 97.2 | 12 |
| B3_HK5671 | chr6 | 46,257,577 | 78 | 80 | 97.5 | 12 |
| B3_HK5671 | chr8 | 122,860,180 | 77 | 79 | 97.5 | 12 |
| B3_HK5671 | chr13 | 74,453,180 | 41 | 42 | 97.6 | 12 |
| B3_AU1054 | chr14 | 26,605,863 | 89 | 91 | 97.8 | 12 |
| B3_AU1054 | chr16 | 46,481,122 | 91 | 93 | 97.8 | 12 |
| B3_AU1054 | chr3 | 166,329,314 | 44 | 45 | 97.8 | 12 |
| B3_HK5671 | chr12 | 67,968,041 | 87 | 89 | 97.8 | 12 |
| B3_AU1054 | chr1 | 87,994,642 | 70 | 71 | 98.6 | 12 |
| B3_HK5671 | chr23 | 136,704,669 | 68 | 69 | 98.6 | 12 |
| B3_HK5671 | chr5 | 128,659,574 | 99 | 100 | 99.0 | 12 |
| B3_HK5671 | chr2 | 143,534,188 | 109 | 110 | 99.1 | 12 |
| B3_HK5671 | chr14 | 26,605,863 | 81 | 81 | 100.0 | 12 |
| B3_HK5671 | chr3 | 166,329,314 | 38 | 38 | 100.0 | 12 |
| B3_HK5671 | chr5 | 166,150,005 | 60 | 60 | 100.0 | 12 |
| B3_HK5671 | chr6 | 163,225,362 | 73 | 73 | 100.0 | 12 |
| B5 | chr20 | 38,282,587 | 78 | 81 | 96.3 | 13 |
| B3_AU1054 | chr14 | 86,373,675 | 60 | 66 | 90.9 | 14 |
| B3_AU1054 | chr15 | 46,555,032 | 55 | 60 | 91.7 | 14 |
| B3_HK5671 | chr15 | 46,555,032 | 76 | 80 | 95.0 | 14 |
| B3_HK5671 | chr14 | 86,373,675 | 72 | 72 | 100.0 | 14 |
